# Long-chain polyphosphates induce glomerular microthrombi and exacerbate LPS-induced acute kidney injury in mouse

**DOI:** 10.1101/2025.07.17.664574

**Authors:** Anniina Pirttiniemi, Hanne Salmenkari, Krishna Adeshara, Jere Lindén, Sanna Lehtonen, Niina Sandholm, Per-Henrik Groop, Markku Lehto

## Abstract

Polyphosphates are evolutionarily conserved anionic polymers mediating pleiotropic functions in eukaryotes and prokaryotes, depending on their chain-length. Bacteria typically synthetize long-chains, while human platelets harbour exclusively medium-chains. Polyphosphate-mediated lung and liver-injury have been reported in experimental mouse models but their effects on the kidney remain undefined. Here we assessed kidney histopathology and cytokine levels following intravenous administration of medium-chain (P100) and long-chain (P700) polyphosphates and their synergistic effects with lipopolysaccharides (LPS) in mice. We found that P700 induced albuminuria, renal *Kim-1* and *Lcn2* transcription, focal renal damage with glomerular microthrombi, tubular degeneration, granular phenotype of slit diaphragm components nephrin and ZO1, and enlarged electron dense vesicles in podocyte cytoplasm indicating lysosome swelling. P700 combined with LPS induced marked multifocal acute tubular necrosis in the cortex, and augmented LPS-induced proinflammatory cytokine levels. No notable effects were seen with P100, indicating that PolyP-mediated kidney injury development is dependent on chain-length. We conclude that long-chain polyphosphates may play a procoagulant role behind kidney injury, by inducing microthrombi characteristic of thrombotic microangiopathy and augmenting cytokine levels under inflammatory conditions.

## Introduction

Polyphosphates (PolyPs) are evolutionarily conserved linear phosphate polymers, which have been studied increasingly in recent years for their ability to regulate various biological processes^1^. They are synthetized ubiquitously in eukaryotes and prokaryotes. Longer chains of 100 to 1000 phosphate residues are synthetized in bacterial cells, where they regulate cellular energy metabolism and virulence^2,3^. Upon lysis of pathogenic gram-negative bacteria, PolyPs may be released together with pathogen-associated molecular patterns (PAMPs), such as lipopolysaccharide (LPS)^4,5^. In rodent tissues PolyPs have been identified in several organelles, e.g. lysosomes, mitochondria, dense granules, and nuclei^6–8^ with chain lengths ranging from 50 to 800 phosphate residues and concentrations ranging from ∼25 to 120 μM^9^. Particularly high concentrations up to 130 mM of medium-chain PolyPs (∼50 to 75 phosphate residues) are stored in platelets and mast cells, from where they can be secreted upon activation^10–12^.

PolyPs have pleiotropic impact on cellular function and previous studies have shown variable outcomes depending on their chain length, dose, and route of administration. Exogenously administered PolyPs have been reported to improve wound healing^13,14^, intestinal health^15–17^, and bone mineralization^18^, while endogenous PolyP accumulation in the brain has been associated with neurodegenerative diseases, such as amyotrophic lateral sclerosis and frontotemporal dementia^19^. PolyPs modulate systemic inflammation in a chain length-dependent manner: long-chain PolyPs (∼700 phosphate residues) can inhibit type I interferon signaling in immune cells, augment *E. coli*/LPS-induced proinflammatory cytokine release and mortality, while shorter PolyPs (∼150 phosphate residues administered intravenously) have conversely showed protective effects against mortality in mice^20–23^. Previous studies have demonstrated that high doses of PolyPs administered intravenously, intraperitoneally or intratracheally can induce acute lung and liver injury in mice, mediated by immunomodulation and coagulation^3,22,24–26^, however the impact of PolyPs on the kidneys has not been assessed.

Acute kidney injury (AKI) is a major cause of morbidity and mortality worldwide and can predispose to conditions such as multiorgan failure and chronic kidney disease. Infection-induced renal injury, such as AKI, may be directly mediated by microorganisms or indirectly mediated by immune system activation. Sepsis-induced AKI is common among the severe cases and is characterized by kidney inflammation, cytokine release, and microcirculatory alterations causing hypoperfusion and hypoxia^27^. AKI has been experimentally modeled in mice most commonly with ischemia–reperfusion injury, nephrotoxic agents, and in models of septic AKI, with cecal ligation puncture or endotoxemia^28,29^.

Here our aim was to investigate the impact of PolyP exposure on the kidney by assessing kidney injury markers, histopathology, and cytokine levels in mice. Since different chain lengths of PolyPs may be released to circulation e.g. from platelets or bacteria^1,5,12^, we studied the effects of intravenously administered medium-chain (P100), and long-chain (P700) PolyPs, as well as their impact on LPS-induced AKI phenotype. We hypothesized that PolyPs can affect kidney function or alter LPS-induced kidney injury development in a chain length-dependent manner.

## Results

### Long-chain polyphosphates induce albuminuria and transcriptional markers of kidney injury and augment LPS-induced cytokine levels

P100 or P700 treatment induced no clinical signs of sickness in the mice during our pilot experiment (3-day follow up, n=2) or the final experiment (20-hour follow up, n=4), whereas LPS-treated mice exhibited typical clinical signs of sepsis within 20 hours in the pilot (n=2) and in the final experiment (n=8): hunched posture, piloerection, and dehydration. In the final experiment P100 did not alter the LPS-induced clinical signs (n=8), whereas P700 worsened them as the condition of the mice in the LPS+P700 group deteriorated rapidly and two out of the eight mice in the group died prior to sample collection at 20 hours post treatment.

Kidney function was assessed 20 hours post treatment in the final experiment by urine albumin-creatinine ratio (uACR). While P100 treatment had no significant effects, P700 elevated uACR significantly to similar levels as LPS, compared to healthy controls. The highest uACR levels were seen in the LPS+P700 group. Of note, P100 or P700 did not elevate urine creatinine levels, as was seen in the LPS-treated groups (Fig. 1a).

**Fig. 1.**
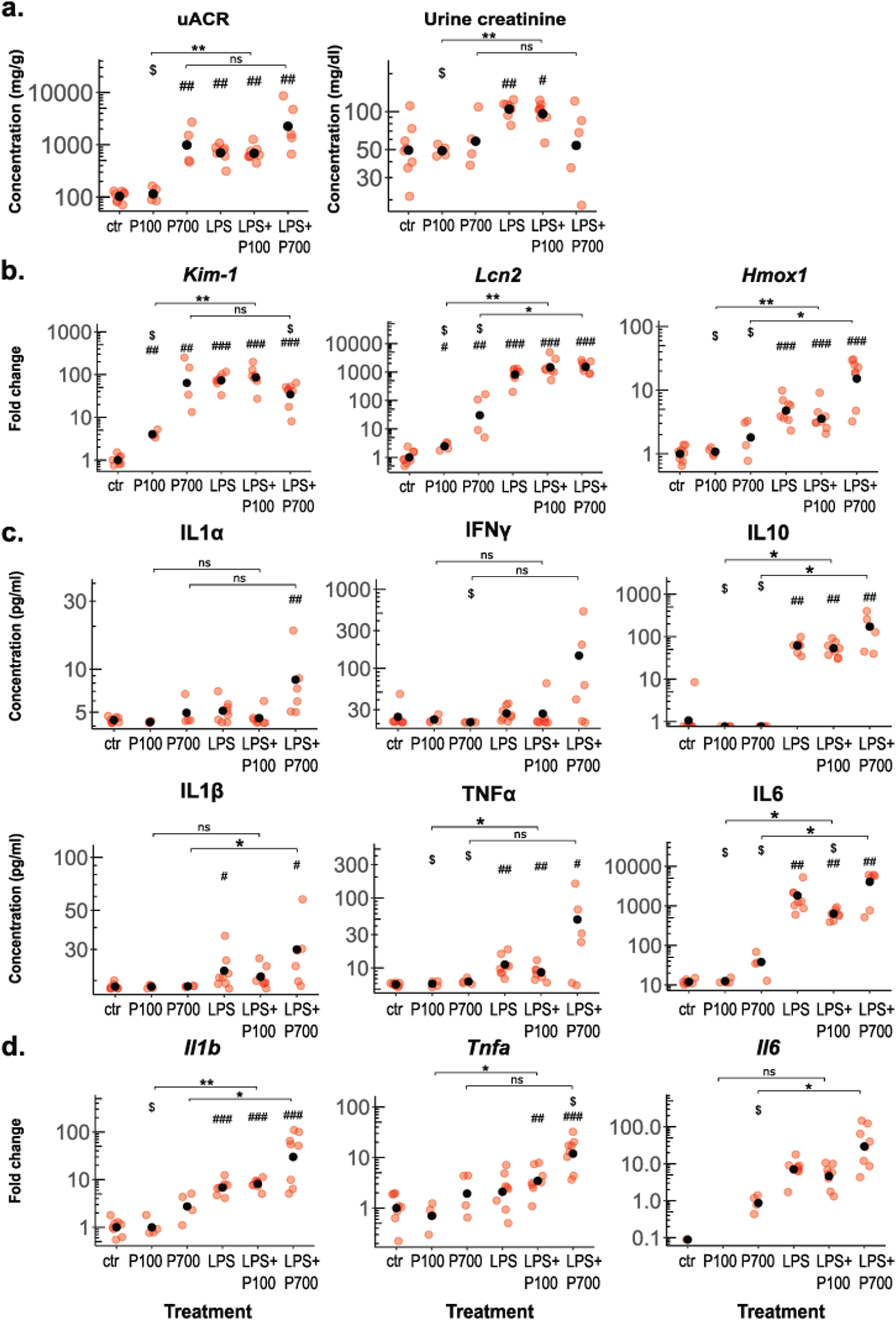
Markers of acute kidney injury in polyphosphate and LPS-treated mice. Mice were treated with medium-chain polyphosphates (P100), long-chain polyphosphates (P700) alone and in combination with LPS (n: 4-8 mice per group). (a) Urine albumin-creatinine ratio (uACR) and urine creatinine levels of the treated mice. (b) Transcriptional levels of kidney injury markers *Kim-1* and *Lcn2,* and *Hmox1*, measured by RT-qPCR from the kidney tissue. (c) Serum cytokine levels of the treated mice measured by a multiplex ELISA assay. (d) Transcriptional levels of cytokines measured by RT-qPCR from the kidney tissue. (a and c) For urine and serum measurements red dots indicate the concentration values from each mouse and the black dots indicate the arithmetic means per group. (b and d) For RT-qPCR measurements red dots indicate the relative gene expression of each mouse compared to control group. Black dots indicate the geometric mean per group, scaled as one in the control group. (d) For or *Il6,* the P700 group is exceptionally scaled as one. Benjamini-Hochberg adjusted significance levels are calculated using the pairwise Mann-Whitney U test., *: p < 0.05, **: p < 0.01, ***: p < 0.001. Symbols used for significance: #: significance compared to the control group, $: significance compared to the LPS-treated group, ns: non-significant.

Transcription of kidney injury markers, *Kim-1, Lcn2*, and *Hmox1,* were analyzed with RT-qPCR. P100-treatment caused a minor elevation to *Kim-1* and *Lcn2,* whereas P700-treatment elevated all injury markers more markedly. LPS significantly elevated all injury markers, and P700 treatment moderately lowered the LPS-induced transcription of *Kim-1* but elevated the LPS-induced transcription of *Hmox1* (Fig. 1b)

Cytokine levels were measured from the serum by multiplex ELISA (Fig. 1c) and from kidney tissue by RT-qPCR (Fig. 1d). While P100 and P700-treatment did not cause significant cytokine elevation compared to controls, LPS caused a significant elevation to serum IL10, IL1β, TNFα and IL6 and transcription of *Il1b* in kidney tissue. The LPS-induced serum levels of IL6 were slightly decreased by P100, whereas P700 caused an increasing trend in all LPS-induced cytokine levels in serum and kidney tissue (Fig. 1c-d).

We assessed the transcription of interferon stimulated genes, *Ifit2*, *Isg15*, and *Oas1a*, which are downregulated by P700 in human and murine leukocytes ^20,21^. Conversely, in the kidney tissue, P100 or P700 did not notably affect the transcription of these genes, but P700 augmented the LPS-induced transcription of *Ifit2*, and *Isg15* (Supplementary Fig. S1a).

### Long-chain polyphosphates induce focal tubular inflammation, apoptosis and cast formation in the kidney, and marked acute tubular necrosis when combined with LPS

Histopathological signs of kidney injury were assessed from H&E and PAS-stained kidney sections (Fig. 2a-b, Supplementary Table S1). While P100-treatment caused no histological alterations, P700-treated mice displayed foci of tubular degeneration and acute tubular necrosis in the cortex and the outer stripe of the outer medulla. The degenerated tubular segments exhibited granular eosinophilic cytoplasm and the necrotic segments, nuclear pyknosis, karyorhexis or –lysis and cell sloughing. Additionally, protein casts were present in the collecting ducts or tubules (pars recta) and the collecting ducts in inner medulla (Fig. 2a, Supplementary Table S1). Glomerular congestion and disruption of adjacent proximal tubule brush borders were observed focally in the P700-treated mice (Fig. 2b). LPS-treated mice typically displayed minimal tubular degeneration in multiple foci in the subcapsular cortex, with epithelial cells showing mild swelling, increased eosinophilia and vacuolated cytoplasm (Fig. 2a). P100 did not modify the LPS-induced histological changes apart from occasional dilated glomerular capillaries (Fig. 2a-b). The histological lesions in the LPS+P700-treated group were similar to P700 and LPS individually, but additionally exhibited marked multifocal acute tubular necrosis (Fig. 2a-b, Supplementary Table S1). Acute phosphate poisoning/nephrotoxicity^30^, was ruled out by a Von Kossa staining, which showed no signs of calcium deposits in the kidney (Supplementary Fig. S1b).

**Fig. 2.**
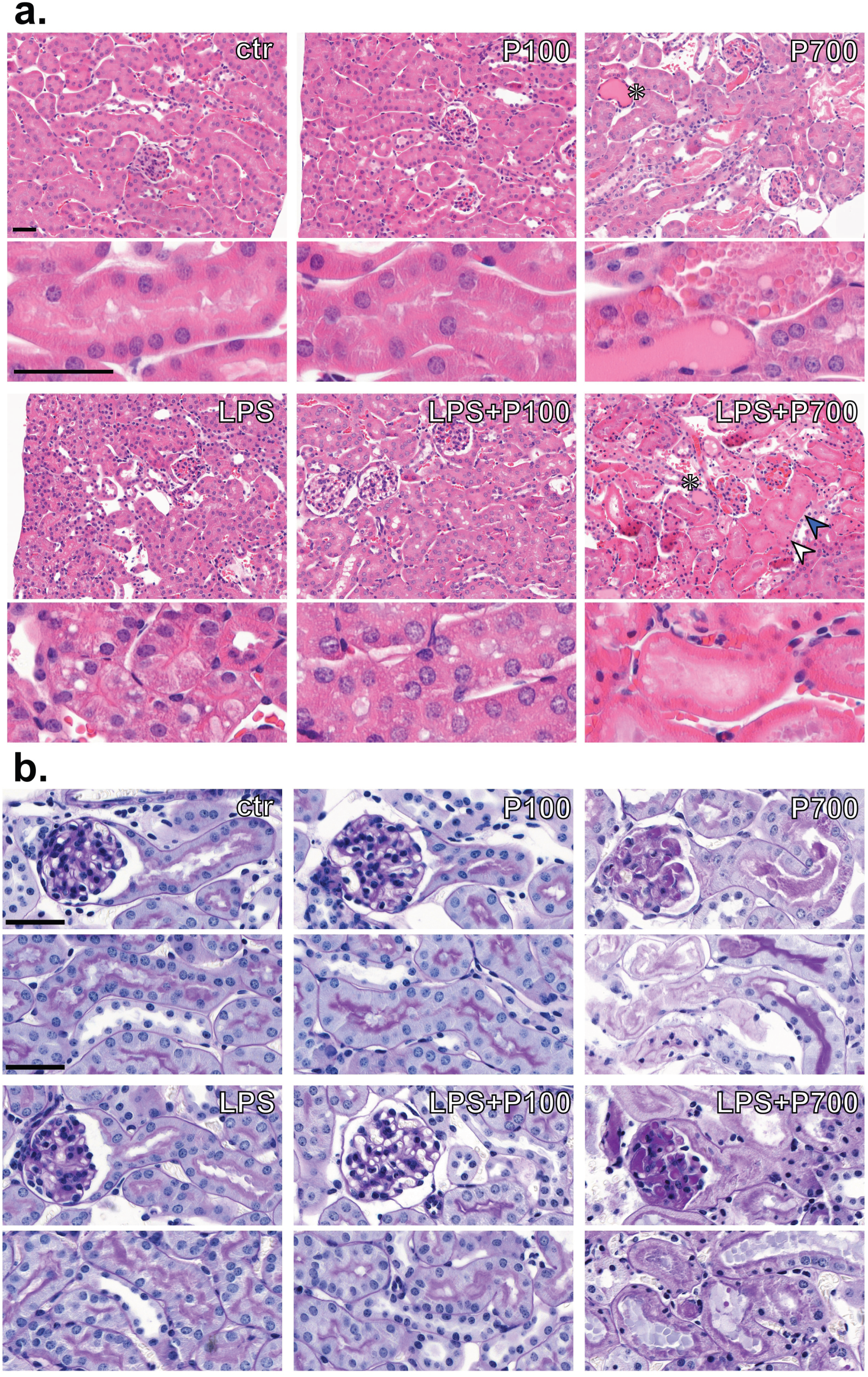
Histopathological manifestations of acute kidney injury seen in polyphosphate and LPS-treated mice. (a) Hematoxylin & Eosin staining showing kidney histopathological changes seen in mice treated with medium-chain polyphosphates (P100), long-chain polyphosphates (P700) alone and in combination with LPS. P700-treated mice show foci of acute tubular necrosis and tubular casts most prominent in the outer medulla (white asterisks). Mice treated with the combination of LPS and P700 additionally exhibit extensive acute tubular necrosis in the cortex, manifested as cytoplasmic eosinophilia, pyknotic nuclei (white arrowhead), karyolysis (blue arrowhead) and tubular casts. LPS and LPS+P100-treated mice display minimal tubular degeneration, with mildly swollen and vacuolated cytoplasm. (b) PAS-staining shows glomerular congestion with deposits in the capillary loops, as well as loss of adjacent proximal tubule brush border integrity in the affected foci in the P700 and LPS+P700-groups. Scale bars 40 µm.

Proximal tubule brush border morphology and tubular apoptosis were further assessed with immunostaining for lotus tetragonolobus lectin (LTL), and cleaved caspase 3 (Fig. 3a). No changes were seen in the P100-group compared to controls, whereas the P700-group showed loss of proximal tubule brush border integrity and tubular apoptosis within the proximal tubuli, focally in the outer medulla. LPS and LPS+P100-treated mice showed only few apoptotic cells in the medulla and cortex. LPS+P700-treated group displayed similar apoptotic proximal tubuli as P700-treated group in the medulla, but additionally exhibited tubular and glomerular cells in active apoptosis in the cortex (Fig. 3a).

**Fig. 3.**
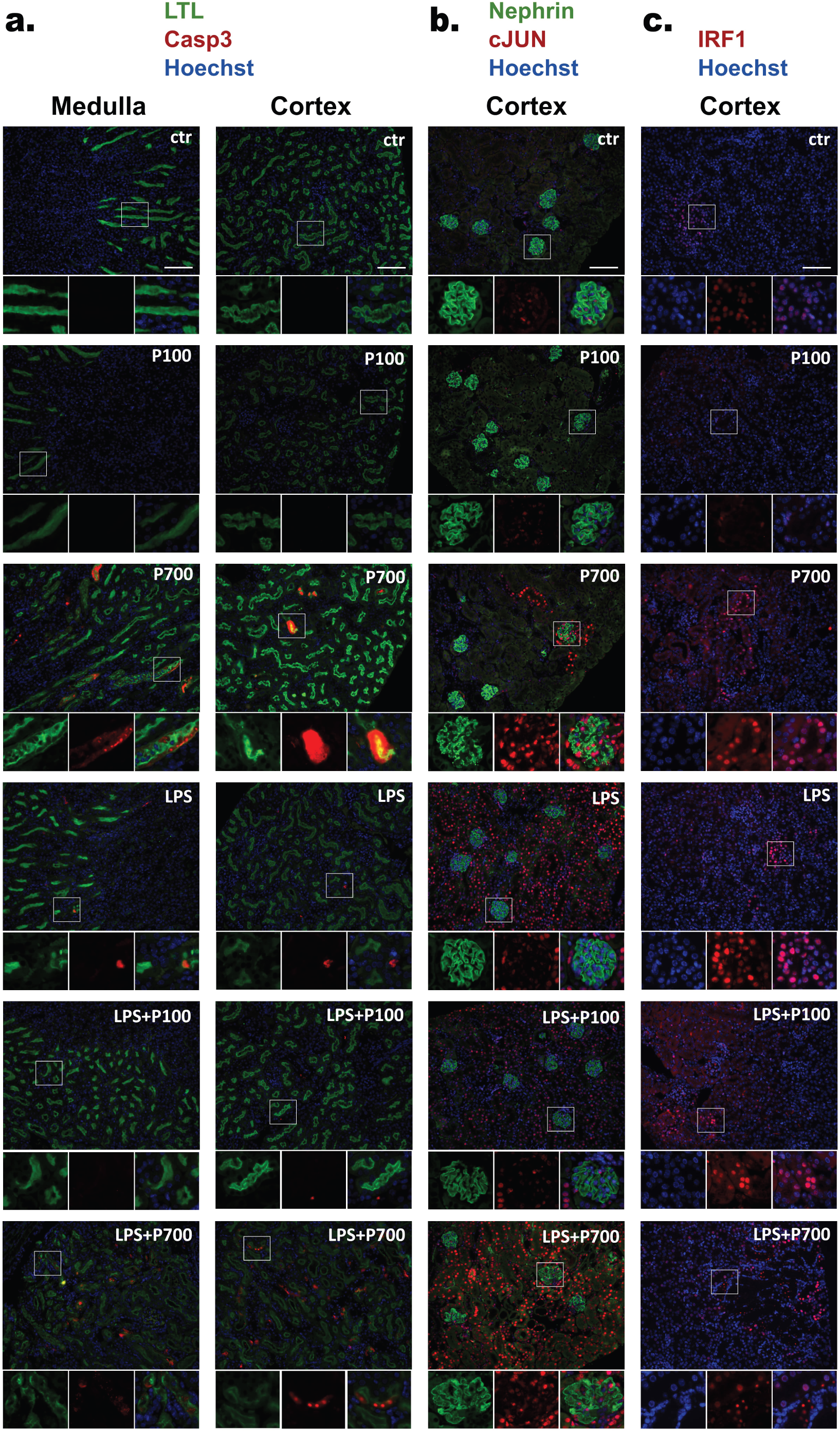
Kidney tubular apoptosis and expression of inflammatory markers cJUN and IRF1 in polyphosphate and LPS-treated mice. Mice were treated with medium-chain polyphosphates (P100), and long-chain polyphosphates (P700) alone and in combination with LPS. (a) Immunofluorescence staining of apoptotic cells (cleaved caspase 3, red) and kidney proximal tubule brush borders (LTL, green) revealed apoptosis inside disrupted proximal tubules in mice treated with P700 most prominently in the outer medulla. In the LPS+P700-treated mice several individual apoptotic tubular and glomerular cells are additionally seen in the affected foci in the cortex. Only few apoptotic cells are seen in the LPS and LPS+P100-treated groups. (b) The kidney cortex showed distinct cJUN-positive (red) foci in P700-treated mice, which were located adjacent to the glomeruli with a granular nephrin (green) phenotype. Mice treated with LPS, LPS+P100, and LPS+P700 showed evenly distributed cJUN expression throughout the cortex, independent of the glomeruli with a granular nephrin phenotype. (c) Foci with IRF1-positive (red) nuclei were slightly increased in the kidney cortex in response to treatment with P700 or LPS compared to controls. Strong IRF1-expression was seen in pyknotic nuclei in necrotic areas of LPS+P700-treated mice. Nuclei stained with Hoechst (blue), scale bar 100 µm.

Immunostaining for cJUN, a marker associated with several kidney diseases^31,32^ showed only few weakly stained nuclei in the glomerular and tubular cells in the controls and P100-treated mice. All P700-treated mice displayed distinct isolated foci with marked glomerular and tubular cJUN-positive nuclei in the cortex, whereas all mice in the LPS-treated groups displayed diffusely distributed tubular cJUN expression in the cortex (Fig. 3b). Staining for the proinflammatory transcription factor IRF1 ^33^ showed cortical foci with tubular nuclear expression in all groups. Although the number of IRF1-positive foci were only slightly increased in the P700 and LPS-treated groups compared to controls, notably strong nuclear IRF1-expression was seen in the apoptotic/necrotic foci in the LPS+P700 group, especially in the pyknotic nuclei, which no longer stain positive for Hoechst (Fig. 3c).

Leukocyte infiltration in the kidney tissue was assessed with immunostaining of a tissue resident macrophage marker, F4/80. While P100 did not cause a general increase in F4/80-covered area compared to controls, all other treatment groups showed an increasing trend for macrophages. The highest macrophage covered areas were seen in the LPS+P700-treated mice, particularly in the most damaged foci (Supplementary Fig. S2a-b). Moreover, glomerular intracapillary leukocytes were seen in transmission electron microscopy (TEM)-analysis only in the LPS-treated groups (Supplementary Fig. S2c, Supplementary Table S2).

### Long-chain polyphosphate-affected foci in the kidney display glomerular microthrombi comprised of adhered platelets, von Willebrand factor, fibrin and collagen fibrils

The P700-induced glomerular congestion was assessed further with Masson’s trichrome staining, which revealed intracapillary deposits indicative of fibrin (red) and collagen (blue), as well as erythrocyte accumulation (Fig. 4a, Supplementary Table S1). Collagen fragments may be released to circulation by matrix metalloproteinase-mediated degradation from sources like exposed endothelial or glomerular basement membrane and participate in early thrombus formation with fibrin^34,35^. These signs of thrombotic events were further supported by Immunostaining with platelet-marker, CD61, and the inductor of platelet adhesion, von Willebrand factor (vWF), which revealed conspicuous glomerular microthrombi. Several glomeruli in the P700-treated group displayed CD61-positive platelet accumulation in the capillary loops and the adjacent arterioles (Fig. 4b) and notable intracapillary vWF-positive deposits (Fig. 4c). The P100-treated mice showed no signs of thrombosis, apart from a few glomeruli with slight increase of CD61 and vWF expression (Fig. 4a-c, Supplementary Table S1). The dose of LPS used here did not induce glomerular thrombosis. LPS+P700-treated mice showed similar glomerular thrombosis as P700-treated mice but showed more pronounced collagen deposition (blue) in Masson’s trichrome staining (Fig. 4a-c, Supplementary Table S1).

**Fig. 4.**
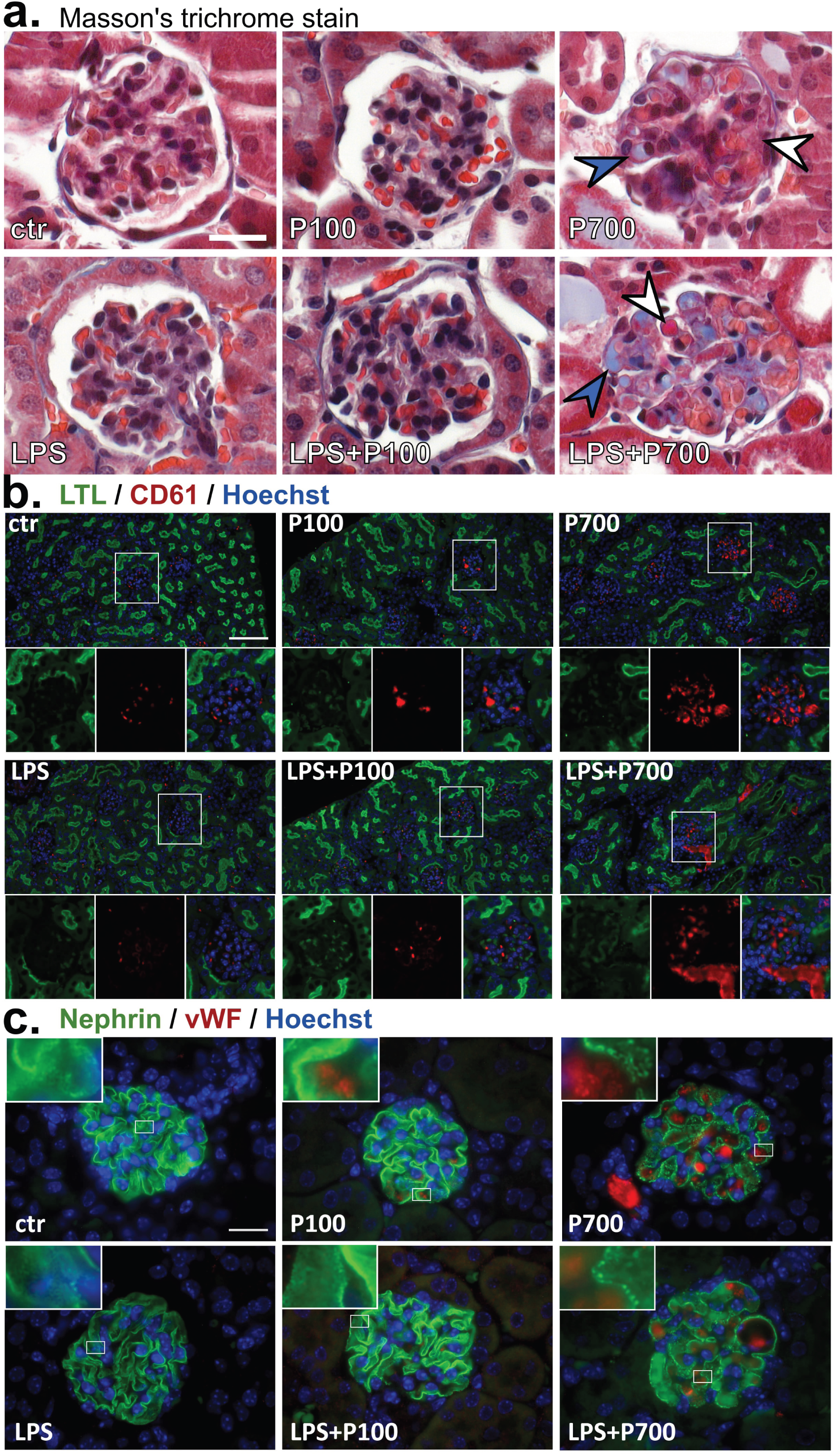
Glomerular microthrombi consisting of fibrin/collagen, von Willebrand factor and platelet accumulation in the kidney cortex of polyphosphate and LPS-treated mice. Mice were treated with medium-chain polyphosphates (P100), long-chain polyphosphates (P700) alone and in combination with LPS. (a) Masson’s trichrome stain shows red stained deposits indicating fibrin (white arrowheads) and blue deposits indicating collagen (blue arrowheads) in the glomerular capillary loops of P700 and LPS+P700-treated mice. Scale bar 20 µm. (b) Immunofluorescence staining of platelets (CD61, red) and kidney proximal tubule brush borders (LTL, green) show platelet accumulation in glomeruli and arterioles of P700 and LPS+P700-treated mice. Nuclei stained with Hoechst (blue), scale bar 100 µm. (c) Immunofluorescence staining showing von Willebrand factor-positive (vWF, red) deposits in the glomerular capillary loops of P700 and LPS+P700-treated mice, with the slit diaphragm marker, nephrin (green) showing a granular phenotype. Nuclei stained with Hoechst (blue), scale bar 20µm.

Thrombic clot composition was confirmed with TEM-imaging of the P700 and LPS-treated groups. Adhered platelets and cellular debris were seen in the capillary loops of the affected glomeruli in the P700-treated mice. LPS-treated mice showed only individual non-adhered intracapillary platelets, similar to controls. The affected glomeruli in the LPS+P700-group displayed structures indicative of collagen fibrils and fibrin strands in addition to adhered platelets and debris (Fig. 5a-b, Supplementary Table S2).

**Fig. 5.**
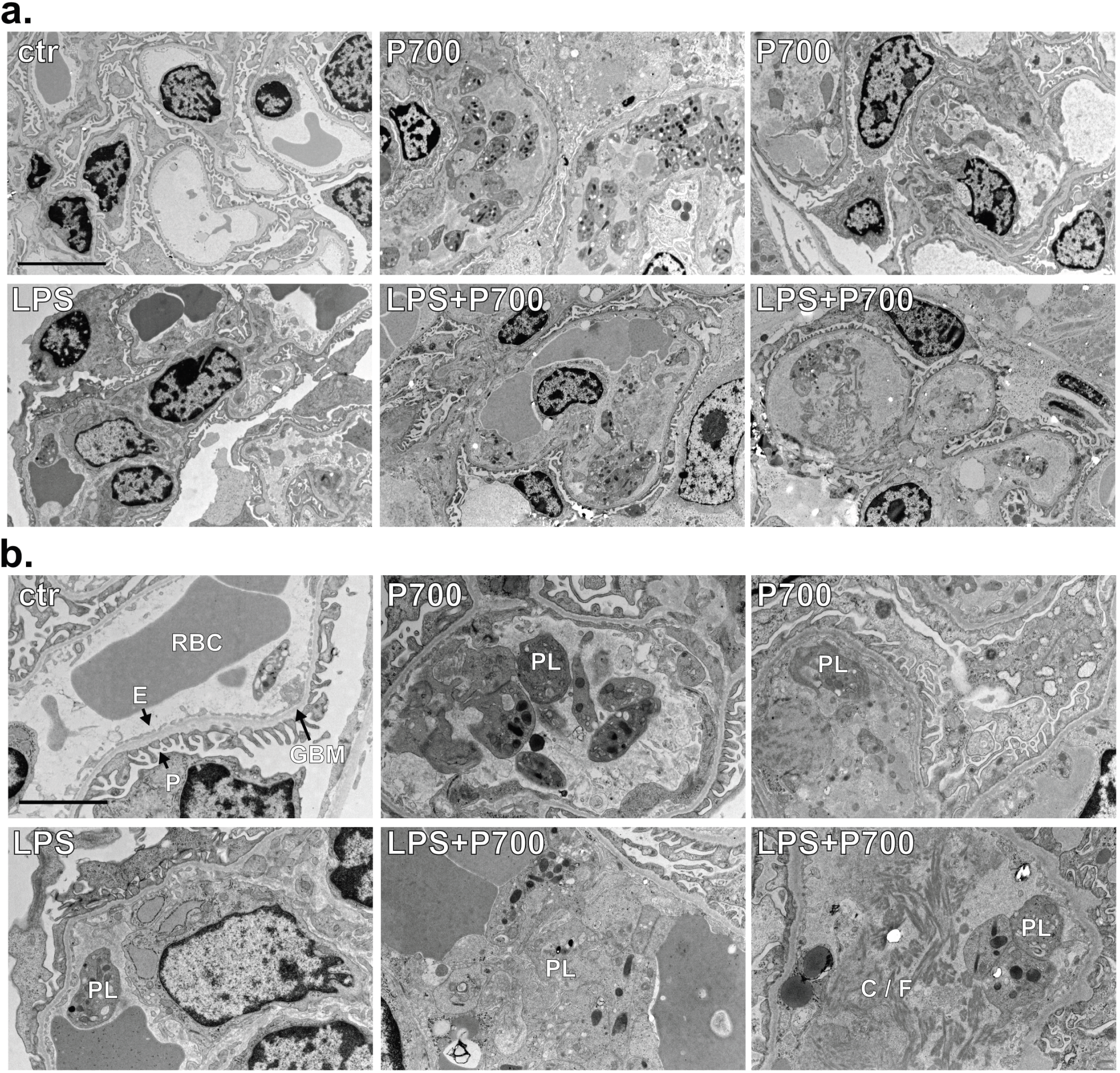
Transmission electron microscopy imaging of the glomerular microthrombi in the kidney cortex of long-chain polyphosphate and LPS-treated mice. Transmission electron microscopy images showing glomerular microthrombi consisting of adhered platelets and cellular debris in P700-treated mice. LPS+P700-treated mice additionally display structures indicative of collagen fibrils / fibrin strands. Individual non-adhered intracapillary platelets can be seen in the control and LPS-treated mice. (a) 1200x magnification, scale bar 5 µm. (b) 3000x magnification, scale bar 2 µm. Fenestrated endothelium (E), podocyte foot processes (P), glomerular basement membrane (GBM), red blood cell (RBC), platelets (PL), collagen fibrils / fibrin strands (C/F).

### Long-chain polyphosphate-affected glomeruli display mislocalization of slit diaphragm proteins and enlarged lysosomes in podocyte cytoplasm

Glomerular slit diaphragm composition was assessed with immunostaining for nephrin and ZO1, which showed an evenly distributed expression lining the glomerular capillary loops in controls (Fig. 6a). A distinct granular phenotype for both proteins, with more distinguishable foot processes, was observed particularly in the P700-treated mice, and a granularity score (ranging from 0 to 5) was used to assess the degree of granularity in all groups (Fig. 6b, Supplementary Fig. S3a). While P100 caused only a slight increase in the granularity scores, P700-treatment induced notable granularity in several glomeruli. LPS induced lower but significant levels of granularity. The LPS+P700 group showed the highest proportion of overall granularity for nephrin, while the P700 group showed the highest overall granularity for ZO1 (Fig. 6a-b, Supplementary Fig. S3a). Nevertheless, the glomerular nephrin and ZO1 granularity scores correlated with each other in the P700, LPS, and LPS+P700 groups and when all groups were combined (Supplementary Fig. S3b). Of note, the granularity was global, affecting the entire glomerulus, although the granular glomeruli were focally distributed in the cortex in all affected groups. Only small fold changes between the treatment groups were seen for transcriptional levels of the nephrin-encoding *Nphs1* and podocalyxin-encoding *Podxl* (Fig. 6c).

**Fig. 6.**
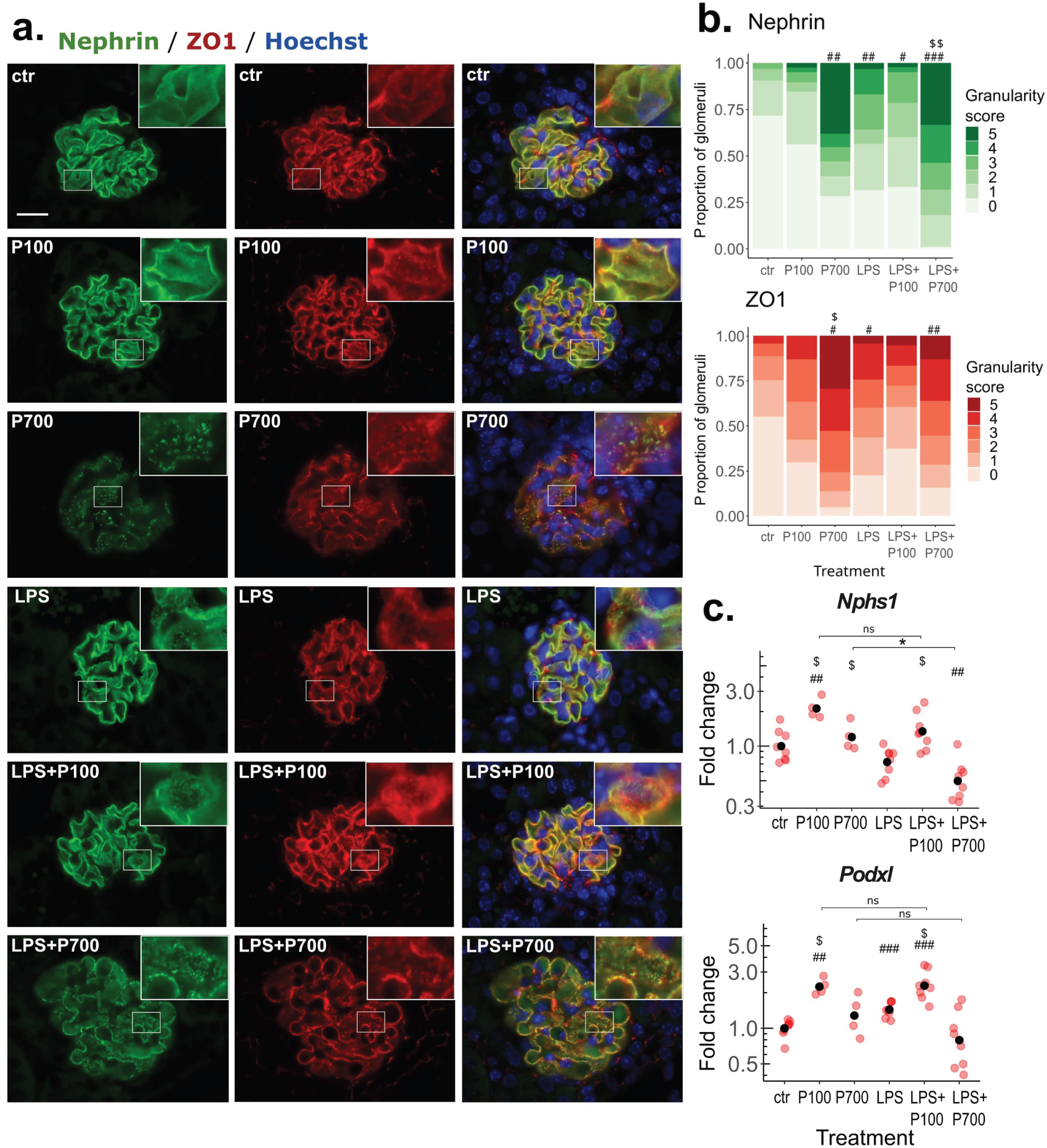
Long-chain polyphosphate-induced granular phenotype of slit diaphragm components, nephrin and ZO1. (a) Representative images of the glomeruli, immunostained with the podocyte-marker, nephrin (green), and tight junction-marker, ZO1 (red), in mice treated with medium-chain polyphosphates (P100), long-chain polyphosphates (P700) alone and in combination with LPS. Nuclei stained with Hoechst (blue), scale bar 20 µm. (b) Proportions and degree of granularity of the nephrin and ZO1-imaged glomeruli. Stacked bars represent the treatment group average of mouse granularity score proportions. Significance levels between the groups are calculated using a granularity score mean value for each mouse (n: 4-8 mice per group, 7-27 imaged glomeruli per mouse). (c) Transcription of nephrin-encoding Nphs1, and podocalyxin-encoding Podxl measured by RT-qPCR from the kidney tissue. Red dots indicate the relative gene expression of each mouse compared to the control group. Black dots indicate the geometric mean per group, scaled as one in the control group. (b-c) Benjamini-Hochberg adjusted significance levels are calculated using the pairwise Mann-Whitney U test., *: p < 0.05, **: p < 0.01, ***: p < 0.001. Symbols used for significance: #: significance compared to the control group, $: significance compared to the LPS-treated group, ns: non-significant.

The connection between slit diaphragm integrity loss and the glomerular microthrombi was investigated by comparing glomerular nephrin granularity score with vWF-positive area per frame. Most significant correlation was seen in the P700 and the LPS+P700 groups, but also in the P100 group, controls, and when all groups were combined. Interestingly, no correlation was seen in the LPS and LPS+P100 groups. (Fig. 4c, Supplementary Fig. S4a-c). Moreover, in the P700-treated mice the glomeruli displaying notable nephrin granularity were generally localized overlapping or adjacent to the foci of cJUN-positive tubuli (Fig. 3b).

Glomerular podocyte and endothelial cell ultrastructural changes were analyzed with TEM in the P700 and LPS-treated groups. P700-treted mice showed enlarged electron-dense vesicles, characteristic of lysosomes in podocyte cytoplasm of the affected glomeruli. While enlarged podocytic vesicles/lysosomes were not seen in LPS-treated mice, LPS+P700-treated mice showed the most notable increase in podocytic vesicle size (Fig. 7a-b, Supplementary Fig. S5). In glomerular endothelial cells, P700 did not cause an increase in cytoplasmic vesicle size, however some increase was seen in the LPS and LPS+P700 groups (Fig. 7a-b, Supplementary Fig. S6). The most severely affected glomerular segments in the P700, LPS, and LPS+P700-treated mice showed also loss of morphology e.g. podocyte vacuolization, foot process fusion and loss of endothelial fenestration (Fig. 7a-b, Supplementary Table S2).

**Fig. 7.**
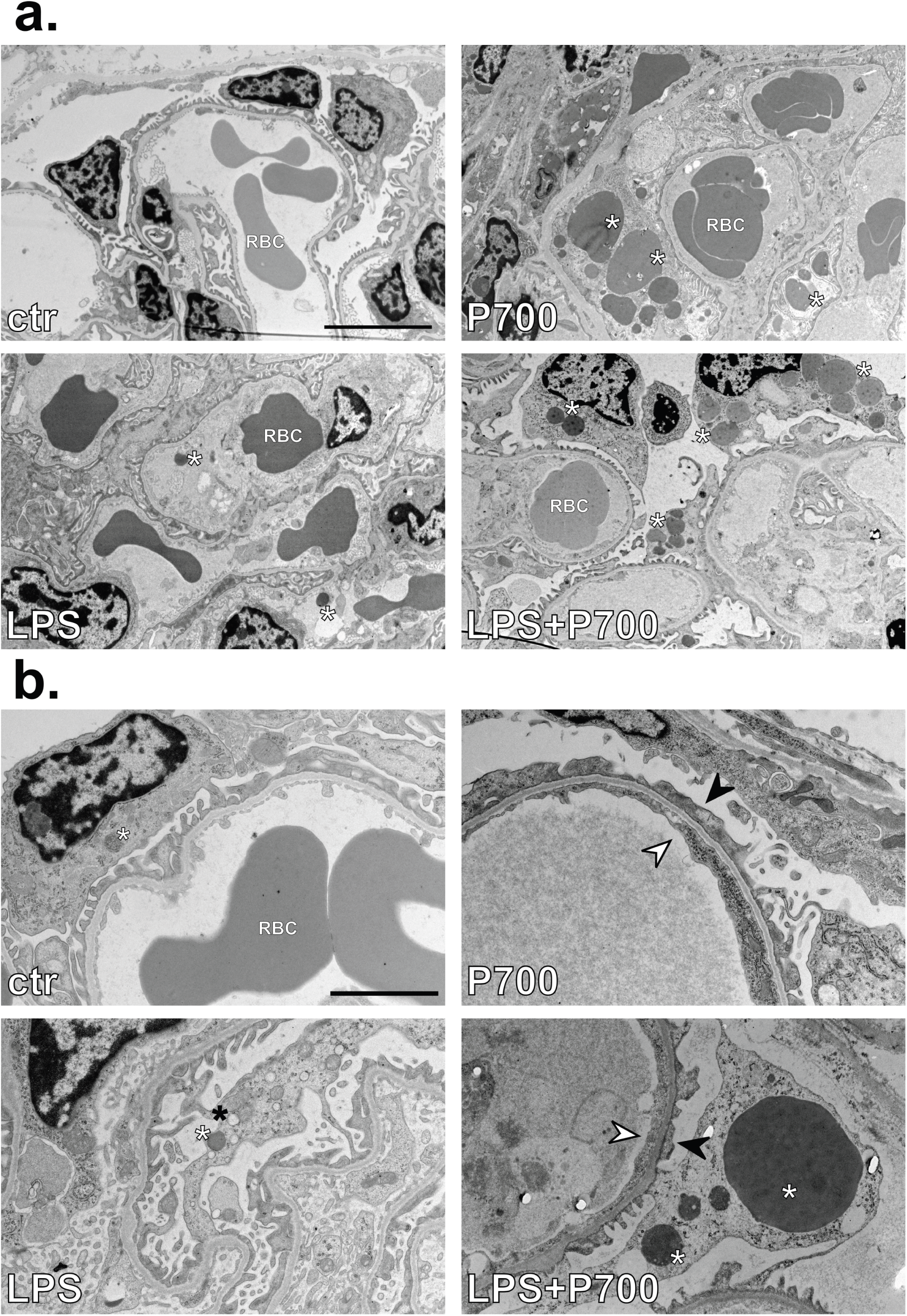
Lysosome swelling and ultrastructural changes in glomerular podocytes and endothelial cells in the mice treated with long-chain polyphosphates and LPS. (a-b) Transmission electron microscopy imaging of glomerular podocytes and endothelial cells in mice treated with long-chain polyphosphates (P700) alone and in combination with LPS, shows large podocytic electron-dense vesicles characteristic of lysosomes (white asterisks) in P700 and LPS+P700-treated mice. (b) Loss of endothelial fenestration (white arrowheads) and podocyte foot process morphology (black arrows) was also seen in the most severely affected glomeruli. Some podocytic and endothelial lysosome enlargement as well as vacuolization (back asterisk) was seen in the most affected glomeruli in LPS-treated mice. (a) 1200x magnification, scale bar 5 µm. (b) 3000x magnification, scale bar 2 µm, red blood cell (RBC).

### Long-chain polyphosphates cause focal alterations in tubular kallikrein-kinin system components, and augment LPS-induced transcription of tissue factor

Since Bradykinin receptor B2 (BDKRB2) and Factor XII (FXII) have been reported to be requisite for PolyP-mediated pulmonary thrombus formation, we investigated related hemostatic mediators, such as kallikrein-kinin system components in the kidney tissue^3,26^. Immunostaining for the tissue kallikrein inhibitor, kallistatin, showed expression within the luminal compartments of the proximal tubules, partly overlapping the bush border, marked by LTL in controls (Fig. 8a). While P100 caused no alterations, P700 caused a clear-cut loss of kallistatin expression within the proximal tubules, which also display a damaged brush border, in distinct foci in the outer medulla. LPS did not alter kallistatin expression compared to controls. LPS+P700-treated mice showed similar changes as the P700-treated mice in the outer medulla. Additionally, the most severely affected cortical foci displayed increased kallistatin expression in the luminal compartments of dilated proximal tubuli, which no longer show an intact proximal tubule brush border or typical overlap of kallistatin and LTL (Fig. 8a). BDKRB2 showed a diffuse tubular expression in the cytoplasm and by the nuclear membrane in the control group, whereas increased receptor expression was seen in all treatment groups in the nucleus and nuclear membrane. Cytoplasmic aggregates of BDKRB2 were seen in the most damaged foci in the P700 and LPS+P700 groups (Fig. 8b). Transcription of kininogen 2-encoding *Kng2* showed only small changes between the groups. Kidney tissue transcription of *F3*, which encodes tissue factor (TF), the initiator of the extrinsic pathway of coagulation, was slightly increased in all treatment groups but was augmented in the LPS+P700-treated mice in an amplificatory manner (Fig. 8c).

**Fig. 8.**
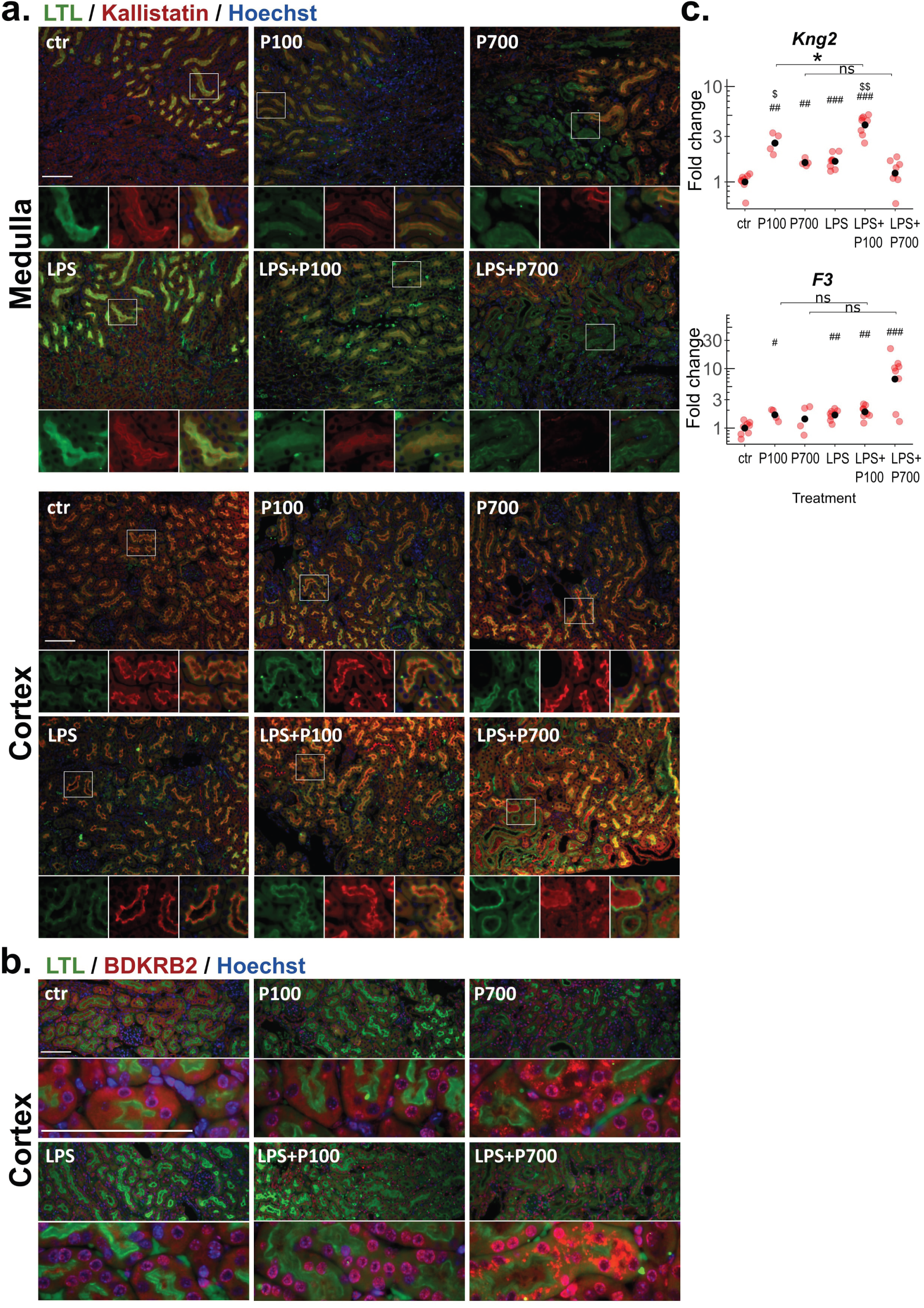
Expression of kallikrein-kinin system components and tissue factor in the kidney of polyphosphate and LPS-treated mice. Mice were treated with medium-chain polyphosphates (P100), long-chain polyphosphates (P700) alone and in combination with LPS. (a) Immunofluorescence staining for tissue kallikrein inhibitor, kallistatin (red) and the proximal tubule marker LTL (green) shows distinct foci in the outer medulla, where both kallistatin expression and the proximal tubule brush border integrity are lost in the P700 and LPS+P700-treated mice. The cortex region of LPS+P700-treated mice displayed augmented kallistatin expression in dilated luminal compartments in the necrotic areas, where proximal tubule brush border integrity is lost. Nuclei stained with Hoechst (blue), scale bar 100 µm. (b) Increased expression of BDKRB2 (red) was seen at tubular cell nuclear membranes in all treatment groups compared to controls. Cytoplasmic aggregation of BDKRB2 was seen in the necrotic areas of P700 and LPS+P700-treated groups. Proximal tubule brush borders stained with LTL (green) and nuclei stained with Hoechst (blue), scale bar 100 µm. (c) Transcription of kininogen 2-encoding *Kng2* and tissue factor-encoding *F3,* measured by RT-qPCR from the kidney tissue. Red dots indicate the relative gene expression of each mouse compared to the control group. Black dots indicate the geometric mean per group, scaled as one in the control group. Benjamini-Hochberg adjusted significance levels are calculated using the pairwise Mann-Whitney U test., *: p < 0.05, **: p < 0.01, ***: p < 0.001. Symbols used for significance: #: significance compared to the control group, $: significance compared to the LPS-treated group, ns: non-significant.

## Discussion

Pleiotropic biological functions have been reported for the evolutionarily conserved PolyP polymers of various chain lengths both in eukaryotes and prokaryotes. As medium or long-chains may be released to circulation e.g. from platelets or bacteria respectively^1,5,12,25,36^, we investigated the impact of intravenously administered medium-chain (P100), and long-chain (P700) PolyPs on kidney histopathology and cytokine levels and assessed the synergistic effects of PolyPs and LPS on AKI development in BALB/c mice. We found that P700 induced glomerular microthrombi and slit diaphragm integrity loss, accompanied by albuminuria and tubular degeneration. Moreover, P700 augmented the LPS-induced AKI and cytokine levels compared to the effects of the individual molecules alone. While LPS-treatment alone induced diffusely distributed cortical histological lesions and upregulation of the inflammatory marker, cJUN, with no signs of glomerular microthrombi, we show that P700-mediated thrombotic damage is focal, likely affecting individual nephrons in the kidney.

In contrast to the pronounced adverse effects of P700 on the kidney, P100 showed only marginal effects in our analysis alone or in combination with LPS. Accordingly, several studies have reported chain length-dependent effects for PolyP exposure, with the longer chains having more capacity to induce coagulation, alter cytokine signaling, complement activity and mortality, but also to facilitate wound healing, and gut epithelial cell function^16,17,20–22,24^. This is perhaps not surprising as millimolar concentrations of shorter chain lengths are endogenous in mammalian platelets’ dense granules, and can be secreted into plasma^12^. PolyPs in the plasma form complexes with divalent cations (namely Ca^2+^) at the surface of cells, such as platelets, where they come into contact with plasma proteins^11,36^. Of note, PolyPs are enzymatically cleaved by phosphatases and have a half-life of ∼2 hours in the plasma *in vitro*^36,37^, although chain length-dependent nanoparticle formation with cations may affect PolyPs solubility and stability^38^. As monophosphate concentration in the blood is maintained at 2.5 – 4.5 mg/dl by excretion/reabsorption by the kidneys^39^, it is plausible that the shorter chain lengths of PolyPs are metabolized and cleared from the circulation faster thereby diminishing their effects. Upon their release into circulation, the long chains may also bind to platelet surfaces and potentiate their thrombotic activity, as membrane-bound PolyPs can activate the contact pathway more efficiently than free soluble PolyPs^40^. As the dose used in our study was based on previous *in vivo* PolyP exposure studies ^21,22^, more studies are needed to clarify serum PolyP concentration and chain lengths in healthy individuals as well as during microbial infections.

In the present study, intravenous P700 exposure induced focal glomerular microthrombi consisting of aggregated platelets, vWF, fibrin and collagen fibrils. The clot composition is characteristic of thrombotic microangiopathy (TMA), which in humans can be caused by e.g. Shiga toxin-producing *E. coli*, complement factor H-autoantibodies (hemolytic uremic syndrome, HUS), sepsis (disseminated intravascular coagulation, DIC) or a deficiency of a vWF-cleaving metalloproteinase, ADAMTS13 (thrombotic thrombocytopenic purpura, TTP)^41,42^. PolyPs participate in thrombosis/coagulation regulation through several mechanisms, such as activation of FXII/contact pathway of coagulation, and factor V, as well as binding and activation of vWF and interference of fibrinolysis^11,26,43–47^. In the context of autoantibody-mediated thrombotic diseases, PolyPs have been shown to modify the conformation of platelet factor 4 (PF4) in a similar manner as other polyanions, heparin (heparin-induced thrombocytopenia, HIT) and DNA (vaccine-induced immune thrombotic thrombocytopenia, VITT), which can eventually lead to platelet-activating autoantibodies^11,48–50^. Endogenous and microbial PolyPs of various lengths may therefore participate in thrombotic diseases through several routes, although autoantibody-mediated effects, such as thrombocytopenia, are not expected in our 20-hour AKI model.

The kallikrein-kinin system regulates hemostasis and inflammation in concert with the renin-angiotensin and the complement systems^51,52^. We found increased nuclear expression of BDKRB2 in the kidney cortical tubular cells in all treatment groups. Knock-out mice for BDKRB2 or FXII were previously reported to be protected from thrombosis and mortality induced by E. coli-derived PolyP (750 mg/kg, i.p.) or platelet-derived PolyP (300 mg/kg, i.v.)^3^. On the other hand, simultaneous knock-out of the constitutively expressed BDKRB2 and the inducible bradykinin receptor B1 (BDKRB1) have been shown to exacerbate AKI in an ischemia reperfusion injury mouse model^53^, as well as diabetic nephropathy development in Akita mice^54^. These studies suggest that BDKRB2 and BDKRB1 may have protective roles through limiting oxidative stress, mitochondrial DNA damage, and expression of fibrogenic genes in the kidney^53,54^. We show here that renal expression of the tissue kallikrein inhibitor, kallistatin, is lost in the disrupted proximal tubule brush border and increased in the dilated lumen in the most severely affected foci in P700 and LPS+P700-treated mice.

Accordingly, kallistatin has been shown to be protective in chronic kidney disease and sepsis-induced tissue injury, though inhibition of oxidative stress, fibrosis, and inflammation^55–57^.

Moreover, we found focally affected glomeruli with granular deposits of two slit diaphragm components: nephrin, a transmembrane junction protein expressed in podocytes, and a tight junction protein, ZO1, in the P700 and LPS+P700-treated mice. The degree of nephrin granularity correlated with the glomerular vWF-positive area in the PolyP-treated groups, indicating a connection between glomerular microthrombi and granular phenotype of the slit diaphragm components. However, some clots may lack vWF or have resolved during our model, as nephrin granularity was seen also without vWF-positive clots in the P700 and LPS+P700 groups. Analysis of ultrastructural changes of the glomeruli revealed large electron-dense vesicles characteristic of lysosomes in the podocyte cytoplasm as a result of P700 or LPS+P700-treatment. The slit diaphragm forms a dynamic zipper-like structure, which participates in the highly selective filtration/retention processes, but the intracellular domain of nephrin is also connected to cellular signaling, mediating podocyte mechanotransduction, cytoskeleton motility, and endosomal vesicle trafficking^58–62^. Interestingly, in Zucker diabetic fatty rats a granular nephrin phenotype has been reported, possibly due to increased expression of its interaction partner PACSIN2, which enhances nephrin trafficking *in vitro*^63^. Our study indicates that the P700-mediated podocyte injury and nephrin granularity may be connected to cellular stress mechanisms such as endocytic recycling and lysosomal degradation following glomerular congestion and thrombus formation. However, further studies are needed to show whether the highly negatively charged P700 has direct effects on the slit diaphragm filtration or lysosome swelling in podocytes.

The P700-treated mice showed varying degrees of tubular damage in conjunction with the foci of affected glomeruli. Notable histological lesions, such as acute tubular necrosis, proximal tubule brush border disruption and apoptosis in the outer medulla were observed in the most severely affected P700-treated mice. However, some albuminuria, nephrin granularity, and cJUN-positive tubular foci were observed in all P700-treated mice, including the mice displaying no clear signs of tubular degeneration. This further supports that the glomerulus may be a primary site of injury by P700 due to thrombus formation leading to glomerular congestion, podocyte injury and albuminuria. Tubular damage is likely to occurs secondary, since proximal tubules and the medulla are often an early site of damage in AKI as the highly metabolically active tubular epithelial cells are sensitive to hypoxia and disturbances in tubular fluid flow and shear stress^64,65^. However, the potential direct effects of P700 on tubular cell function still need to be elucidated.

While P700 affected only isolated foci in the kidney cortex and outer medulla, LPS caused more diffusely distributed histological lesions and expression of cJUN in the cortex. In the P700-treated mice the glomerular nephrin granularity correlated with vWF-positive area and coincided with tubular cJUN expression, but the LPS-treated mice did not show similar correlation or any signs of thrombosis in general, indicating that LPS induces the observed milder degrees of nephrin granularity through a different mechanism. Assessment of the synergistic effects of PolyPs and LPS showed that histologically detectable acute tubular necrosis, albuminuria, *Lcn2* transcription, and cytokine levels were exacerbated in LPS+P700 treatment compared to each molecule alone. Immunothrombosis is a concept describing the connection between coagulation and the innate immune system, which operates though the TF-mediated extrinsic pathway of coagulation in an amplificatory manner^66^. P700-treatment alone increased the TF-encoding *F3*, to similar levels as LPS-treatment, but did not cause cytokine release or clinical signs of sepsis in the mice, as was seen with LPS-treatment. This highlights the crucial role of initial pattern recognition receptor activation and the innate immune system activation before the TF-mediated amplificatory immunothrombosis can occur^66^. While TF is best known for amplifying the coagulation process under pathological conditions^66–68^, the function of TF is bidirectional, as it can also amplify cytokine signaling, through e.g. protease-activated receptors (PARs)^69^, inflammasome activation, and neutrophil extracellular traps (NETs)^70^. PolyPs have also been connected to the induction of NETs^71^, which as negatively charged chromatin structures can further activate the contact pathway and function as a scaffold for platelets, red blood cells, extracellular vesicles, vWF, and TF^66,70^. Comprehensive understanding of various PolyP-associated hemostatic amplificatory mediators still requires extensive investigation *in vivo*.

Our study highlights the importance of thorough mechanistic characterization of individual players behind kidney damage, considering the diversity of kidney injury manifestations seen in animal models and kidney injury types seen in humans. However, we were limited by small urine and blood sample volumes in our study and with the acquired samples could not extensively study e.g. plasma coagulation factors, changes in leukocyte and platelet function, or urine and serum markers in the mice. Urine and serum protein markers, such as NGAL (early-stage marker for glomerular and tubular injury) and KIM-1 (marker for proximal tubule injury), have been proposed to better describe the severity, prognosis, and the specific site of injury in the kidney in humans^27,72–75^. Interestingly, in mouse kidney tissue we found the NGAL-encoding *Lcn2* to be notably more sensitive to LPS than P700, while *Kim-1* transcription responded to LPS and P700 similarly.

Sufficient treatment options for sepsis-related mortality and organ damage are still lacking and anticoagulative TF or FVII inhibitors have not been successful in human trials despite promising preclinical trials in animals^68,76–79^. Further studies are needed to map out treatment options, which better consider the timing and type of AKI and improve kidney function after injury has already occurred. This may include better management of blood coagulation and inflammation^80^. In conclusion, P700-treatment may be used as an effective experimental model to study systemic effects of renal thrombosis, characteristic of TMA (commonly modeled with Shiga toxin, LPS or ADAMTS13 depletion in animals)^42,81^, while in combination with LPS the proinflammatory phenotype is augmented in a manner characteristic of immunothrombosis^66^. PolyPs serve also as interesting targets of treatment in humans, although how much PolyPs contribute to thrombosis in the context of bacterial infections or tissue damage in humans remains to be elucidated.

## Materials and methods

### Experimental animals and procedures

The study was conducted adhering to the ethical guidelines and the ethical permit has been approved by the Finnish Project Authorisation Board (ELLA, ESAVI/6504/2020). The study was conducted using 9-week-old wild-type male BALB/c mice (BALB/cAnNCrl; Scanbur, Karlsunde, Denmark) housed in pairs in individually ventilated cages with environmental enrichment, 12:12 h light-dark cycle, and free access to water and chow. AKI was induced by 5 mg/kg intraperitoneal injection of LPS (from *E. coli* strain O111:B4, Sigma Aldrich, Germany). Based on previous PolyP exposure studies^21,22^ a dose of 10.2 mg/kg (equivalent of 100 µmol/kg of total phosphate residues) of medium-chain PolyPs, P100 (Kerafast, USA, heterogenous size distribution of 40 – 160 phosphate units) or long-chain PolyPs, P700 (Kerafast, USA, heterogenous size distribution of 99 – 1298 phosphate units) were administered by intravenous injection under isoflurane anaesthesia. As requested by ELLA, the used LPS and PolyP concentrations were tested on a smaller number of mice (n=2-3 per goup) in a pilot experiment (24-hour follow-up for LPS-treatment and 3-day follow-up for P100 and P700-treatment). Based on the albuminuria levels in the pilot experiment (data not shown), the number of mice per group in the final experiment was kept to a minimum, with which reliable results were attainable.

The treatment groups in the final experiment were: untreated control (n=8), P100-treated (n=4), P700-treated (n=4), LPS-treated (n=8), LPS+P100-treated (n=8), and LPS+P700-treated (n=8). The mice were sacrificed 20 hours after injection, and the kidneys, sera and urine were collected for subsequent analysis. Two mice in the LPS+P700 treatment group in the final experiment unexpectedly died prior to sample collection and these mice were not included in subsequent analysis of urine or serum samples, but kidney tissue samples were included in further analysis.

### Urine albumin and creatinine measurement

Spot urine was collected prior to sacrificing the mice, however urine was not attainable for three mice in the LPS+P700 treatment group and one mouse in the LPS treatment group due to anuria. Urinary albumin concentrations were measured with ELISA (Bethyl Laboratories, E99-134) and creatinine was measured using a Creatinine Urinary Detection Kit (ThermoFisher, EIACUN).

### Serum sample preparation and Quansys multiplex cytokine assay

Blood samples were collected during terminal anesthesia by cardiac puncture. 0.5 – 1 ml blood was pipetted into serum separator tubes and the tube inverted to dissolve the clotting activator. The tubes were incubated for 30 min at RT before centrifuging at 10 000 g for 5 min. Serum cytokine levels were assayed with the Q-Plex Mouse Cytokine Panel 1, 6-Plex (Quansys Biosciences). Serum samples were diluted 1:2 into the kit sample diluent and the plate was measured with a chemiluminescent reader (Q-view imager LC, Quansys Biosciences).

### Kidney tissue sample preparation for histological staining

The dissected kidneys were cut in sagittal and horizontal plane into four pieces for subsequent processing. For histological analysis, the tissues were fixed with 10% neutral-buffered formalin for 24 h, after which they were stored in 70% ethanol at +4 °C from 24 h to 72 h before tissue processing, paraffin embedding, and sectioning at the Finnish Centre for Laboratory Animal Pathology (FCLAP).

### Histochemical staining and digital slide scanning of kidney sections

Hematoxylin & Eosin (H&E), Periodic acid–Schiff (PAS), Masson’s trichrome stain, and Von Kossa staining for the paraffin sections from all 40 mice were performed by the Finnish Centre for Laboratory Animal Pathology (FCLAP). Whole slide scan images (40 x air objective) of the H&E, PAS, and Masson’s trichrome stained microscopy slides were generated using 3DHISTECH Pannoramic 250 FLASH II digital slide scanner at Genome Biology Unit supported by HiLIFE and the Faculty of Medicine, University of Helsinki, and Biocenter Finland and cropped images acquired and adjusted with SlideViewer 2.6.0 (3DHISTECH).

### Immunohistochemical staining of kidney sections and quantification

Paraffin-embedded 4 µm tissue sections were deparaffinized/rehydrated with a standard xylene-ethanol series. Antigen retrieval was conducted with 20 min gentle boiling in 10 mM Tris, 1 mM EDTA, pH 9 + 0,05% Tween in phosphate-buffered saline (PBS). Once cooled the slides were rinsed with PBS and permeabilized/blocked for 1 hour at room temperature (RT) in 5 % fetal bovine serum (FBS, Thermo Fisher), 0,1% Triton X-100 (Sigma Aldrich) in PBS. The primary antibodies (Supplementary Table S3) were incubated over night at +4 °C in 1% FBS in PBS, after which they were washed three times for 15 min in the same buffer. The secondary antibodies (anti-Rabbit IgG Alexa Fluor 594 Cat. A-21207 and anti-Guinea Pig IgG Alexa Fluor 488 Cat. A-11073, diluted 1:500) or fluorescein-conjugated Lotus Tetragonolobus Lectin (LTL) were incubated for 1 h at RT before the slides were washed three times for 15 min. Nuclear stain with Hoechst (Thermo Fisher, Cat. H1398) was included with the secondary antibody incubation. Autofluorescence quenching was conducted with TrueView (Vector Laboratories, Cat. SP-8400) before the slides were mounted with VectaShield Vibrance mounting medium (Vector Laboratories, Cat. H-1700). Images were acquired with Axio Imager M2 microscope, equipped with Axiocam 503 (Carl Zeiss Microscopy GmbH, Oberkochen, Germany) using Zen 3.1 Imaging software.

For nephrin and ZO1 stained glomeruli the degree of granularity was assessed by scoring the blinded 1000x magnification images on a scale from 0-5 of granularity (4-8 mice per group, 7-27 imaged glomeruli per mouse, 444 glomeruli in total). For von Willebrand factor (3-4 mice per group, 6-11 1000x-imaged glomeruli per mouse, 147 glomeruli in total) and F4/80 (4-8 mice per group, 3-9 200x-imaged frames per mouse) stained kidney sections, cell covered area per frame was quantified with Fiji ImageJ^82^. For all other immunostained markers assessing qualitative changes at least three 200x frames were included from 3-4 mice per group.

### Transmission electron microscopy sample preparation and analysis

For transmission electron microscopy analysis of glomeruli, 1 to 2 mm cubes from the kidney cortex were fixed in 2.5% glutaraldehyde, 2.5% paraformaldehyde, 0.1 M phosphate buffer, pH 7.4 for three hours. The samples were further processed by the Electron Microscopy Unit, HiLIFE, Institute of Biotechnology, (University of Helsinki and Biocenter Finland) as described earlier^83^ and imaged with JEM-1400PLUS (JEOL Ltd, Japan) with an Orius SC 1000B bottom-mounted charge-coupled device camera (Gatan). Glomeruli from treatment groups: controls, P700, LPS, and LPS+P700, were imaged and assessed at 1200x and 3000x magnification. The diameter of electron dense vesicles characteristic of lysosomes, multivesicular bodies or peroxisomes^84^ was quantified from 1200x magnification images using Fiji ImageJ ^82^ (3 mice per group, 3-7 imaged glomeruli per mouse, 1-5 frames per glomerulus).

### RT-qPCR

Kidney tissue was lysed in a bead homogenizer (Analytik Jena) and RNA was extracted using NucleoSpin RNA kit (Macherey-Nagel). 1 µg of RNA was reverse transcribed into cDNA with iScript cDNA Synthesis Kit (Bio-Rad) and further diluted 1:10 in molecular biology grade water. For quantitative reverse transcription PCR (RT-qPCR) 2 µl of template and 0.25 µM of forward and reverse primers together with iTaq Universal SYBR Green Supermix (Bio-Rad) were used in a total PCR reaction volume of 10 µl. Hard-Shell 384-Well PCR Plates (Bio-Rad) were used with the CFX384 Touch Real-Time PCR Detection System thermal cycler (Bio-Rad) with the program: initial denaturation at 95 °C for 30 s followed by 40 cycles of denaturation at 95 °C for 5 s, and annealing at 60 °C for 30 s. Fold changes in gene expression were analyzed based on previously described methods^85,86^ and normalized to housekeeping genes *Eef2* and *S18*. Primer sequences with original references are presented in Supplementary Table S4. New primers were generated using Primer3web version 4.1.0 and Primer-BLAST.

## Statistical analysis

R version 4.2.3 was used for statistical analysis. Pairwise Mann–Whitney U test with Benjamini-Hochberg adjusted p-values was used for 2-group comparisons and Spearman’s rank correlation coefficient was used to assess correlation between glomerular markers.

## Acknowledgments

We thank Heli Krigsman for extensive laboratory analysis, Emma Dahlström, and Stefan Mutter for statistical advice, Mervi Lindman from the Electron Microscopy Unit, HiLIFE, Institute of Biotechnology, (University of Helsinki and Biocenter Finland), and the staff of the Finnish centre for laboratory animal pathology (FCLAP) and Tissue Preparation and Histochemistry Unit (TPHU), Helsinki Institute of Life Science, University of Helsinki. Part of the work was carried out with the support of HiLIFE Laboratory Animal Center Core Facility, the University of Helsinki. Graphical abstract illustrations created with BioRender.com. Parts of this study have previously been presented at the European Association for the Study of Diabetes (EASD) meeting (Stockholm, Sweden): Pirttiniemi A, *et al.* Long-chain polyphosphates cause a distinct kidney injury phenotype and exacerbate LPS-induced acute kidney injury in mouse. Diabetologia (2022) 65 (Suppl 1):S1–S469.

## Data and resource availability

Any additional information or original data (e.g. high-resolution image files) required to reanalyze the data reported in this paper may be acquired from the FinnDiane Study Group/the corresponding author for noncommercial research purposes upon reasonable request.

## Author Contributions

Conceptualization (AP, ML), study design (AP, HS, SL, ML), animal experiments (HS, KA, ML), laboratory data acquisition (AP, HS) data analysis (AP), interpretation of data (AP, HS, KA, JL, SL, NS, PHG, ML), histopathological analysis (AP, JL) preparation of the manuscript (AP). All authors have critically reviewed and approved the final version of the manuscript.

