## Supplementary material for "Long-chain polyphosphates induce glomerular microthrombi and exacerbate LPS-induced acute kidney injury in mouse"

### Table S1. Histopathological changes seen in H&E, PAS, and Masson's Trichrome stained sections.

| Treatment group | Mouse ID | Tubular necrosis/ degeneration | Glomerular thrombus formation | Pathological changes |
|---|---|---|---|---|
| ctr | 1 | 0 | 0 | |
| ctr | 2 | 0 | 0 | |
| ctr | 3 | 0 | 0 | |
| ctr | 4 | 0 | 0 | |
| ctr | 5 | 0 | 0 | |
| ctr | 6 | 0 | 0 | |
| ctr | 7 | 0 | 0 | |
| ctr | 8 | 0 | 0 | |
| LPS | 9 | 1 | 0 | See mouse 13. |
| LPS | 10 | 1 | 0 | See mouse 13. |
| LPS | 11 | 1 | 0 | See mouse 13. |
| LPS | 12 | 1 | 0 | See mouse 13. |
| LPS | 13 | 1 | 0 | Minimal tubular degeneration in multiple foci in the subcapsular cortex. The degenerated tubule cells are mildly swollen and show increased eosinophilia and foamy or vacuolated cytoplasm. |
| LPS | 14 | 1 | 0 | See mouse 13. |
| LPS | 15 | 1 | 0 | See mouse 13. |
| LPS | 16 | 1 | 0 | See mouse 13. |
| LPS+P700 | 17 | 1 | 0 | See mouse 13. + A focal acute coagulative necrosis with some hemorrhages and neutrophil infiltration. Oblong focal necrosis extends from OSOM to cortex. Not present in PAS- or MT-stained slides. |
| LPS+P700 | 18 | 3 | 1 | See mouse 21. Glomerular capillary congestion impedes detection of the thrombi. |
| LPS+P700 | 19 | 1 | 0 | See mouse 13. |
| LPS+P700 | 20 | 1 | 0 | Minimal tubular degeneration in multiple small subcapsular foci. The degenerated tubular epithelial cells show foamy, granular or vacuolated cytoplasm (See 13). Necrotic, exfoliated cells in one tubular segment in the OSOM in one HE section - focal change. |
| LPS+P700 | 21 | 3 | 2 | Moderate / marked acute tubular necrosis. Multifocal cortical, subcapsular and single OSOM segments of tubules showing granular brightly eosinophilic cytoplasm, karyorhexis or -lysis and |

| | | | | |
|---|---|---|---|---|
| | | | | cell sloughing. Tubular protein casts. Eosinophilic casts / globules (fibrin thrombi) and erythrocyte aggregates in the glomerular capillary loops in several glomeruli. |
| LPS+P700 | 22 | 4 | 2 | Marked tubular necrosis. Multifocal tubular necrosis extends to ISOM and inner medulla. |
| LPS+P700 | 23 | 3 | 1 | See mouse 21. Glomerular congestion impedes cast assessment. Marked glomerular congestion. |
| LPS+P700 | 24 | 3 | 1 | See mouse 21. Little medulla present in the section (only some OSOM). Marked glomerular congestion. |
| P700 | 25 | 0 | 0 | |
| P700 | 26 | 2 | 2 | Mild tubular degeneration and necrosis. Some cortical, subcapsular and OSOM tubular segments showing granular brightly eosinophilic cytoplasm, karyorhexis or -lysis and cell sloughing or degenerative changes. Protein casts in the collecting ducts or tubules (pars recta) and the collecting ducts in inner medulla. Glomerular congestion. Eosinophilic casts / globules (fibrin thrombi) and erythrocyte aggregates in the glomerular capillary loops in several glomeruli. |
| P700 | 27 | 1 | 1 | Single deep cortical or OSOM degenerated tubular segments (few necrotic cells). See mouse 13. |
| P700 | 28 | 2 | 1 | See mouse 26. Eosinophilic casts / globules (fibrin thrombi) and erythrocyte aggregates in the glomerular capillary loops in some glomeruli. |
| LPS+P100 | 29 | 1 | 0 | See mouse 13. |
| LPS+P100 | 30 | 0 | 0 | |
| LPS+P100 | 31 | 0 | 0 | |
| LPS+P100 | 32 | 0 | 0 | Focal tubular degeneration in the OSOM in HE. Not present in PAS- or MT-stained slides. |
| LPS+P100 | 33 | 1 | 0 | See mouse 13. Dilated glomerular capillaries. |
| LPS+P100 | 34 | 1 | 0 | See mouse 13. |
| LPS+P100 | 35 | 1 | 0 | See mouse 13. |
| LPS+P100 | 36 | 0 | 0 | |
| P100 | 37 | 0 | 0 | |
| P100 | 38 | 0 | 0 | |
| P100 | 39 | 0 | 0 | |
| P100 | 40 | 0 | 0 | |

Note: Mice 23 and 24 were found dead before sample collection. Hematoxylin & Eosin staining (HE), Periodic Acid-Schiff (PAS), Masson's Trichrome staining (MT), outer stripe of the outer medulla (OSOM), inner stripe of the outer medulla (ISOM). Grading of tubular necrosis/degeneration and glomerular thrombus formation; 0: no change, 1: minimal, 2: mild, 3: moderate, 4: marked

**Table S2. Number of affected glomeruli with observed intracapillary clots, leukocytes, and podocyte vacuolization in transmission electron microscopy analysis.**

| Treatment group | Mouse ID | Adhered platelets / platelet clots | Fibrin strans / collagen fibrils | Intracapillary leukocytes | Podocyte vacuolization | Assessed glomeruli in total per mouse |
|---|---|---|---|---|---|---|
| ctr | 1 | 0 | 0 | 0 | 0 | 3 |
| | 2 | 0 | 0 | 0 | 0 | 6 |
| | 4 | 0 | 0 | 0 | 0 | 6 |
| P700 | 25 | 0 | 0 | 0 | 0 | 7 |
| | 26 | 2 | 0 | 0 | 2 | 3 |
| | 28 | 2 | 0 | 0 | 0 | 6 |
| LPS | 12 | 0 | 0 | 1 | 0 | 4 |
| | 14 | 0 | 0 | 4 | 0 | 7 |
| | 15 | 0 | 0 | 1 | 2 | 4 |
| LPS+P700 | 17 | 0 | 0 | 0 | 0 | 3 |
| | 21 | 3 | 0 | 0 | 0 | 6 |
| | 22 | 4 | 3 | 1 | 1 | 5 |

**Table S3. Primary antibodies used in immunohistochemical staining.** **Target Host**

| Target | Host Animal | Manufacturer | Catalogue number | Used dilution |
|---|---|---|---|---|
| Proximal tubule - Lotus Tetragonolobus Lectin (LTL) | - | Vector Laboratories | #FL-1321 | 1:200 |
| Cleaved caspase-3 | Rabbit | Cell Signaling Technology | #9661 | 1:200 |
| Integrin β3 (CD61) | Rabbit | Cell Signaling Technology | #13166 | 1:250 |
| Von Willebrand factor (vWF) | Rabbit | Novus Biologicals | # NB600-586 | 1:1000 |
| Nephrin | Guinea pig | Progen | #GP-N2 | 1:200 |
| Zona Occludens 1 (ZO1) | Rabbit | Abcam | #ab221546 | 1:1000 |
| Bradykinin Receptor B2 (BDKRB2) | Rabbit | Novus Biologicals | #NBP1-46328 | 1:200 |
| Kallistatin | Rabbit | ThermoFisher | #PA5-96636 | 1:100 |
| cJUN | Rabbit | Cell Signaling Technology | #9165 | 1:200 |
| Interferon regulatory factor 1 (IRF1) | Rabbit | Cell Signaling Technology | #8478 | 1:100 |
| CD45 | Rabbit | Cell Signaling Technology | #70257 | 1:150 |
| F4/80 | Rabbit | Cell Signaling Technology | #70076 | 1:150 |

### Table S4. Primer sequences used in RT-qPCR.

| Gene name | Forward primer | Reverse primer | Reference |
|---|---|---|---|
| *Kim-1* | 5′-TTGCCTTCCGTGTCTCTAAG-3′ | 5′-AGATGTTGTCTTCAGCTCGG-3′ | S1 |
| *Lcn2* | 5′- CCACCACGGACTACAACCAG -3′ | 5′- AGCTCCTTGGTTCTTCCATACAG -3′ | - |
| *Hmox1* | 5′-CACATCCAAGCCGAGAATGC-3′ | 5′-AAGGAAGCCATCACCAGCTTAAA-3′ | - |
| *Il1b* | 5′- CTCCAGCCAAGCTTCCTTGT-3′ | 5′- TCATCACTGTCAAAAGGTGGCA -3′ | S2 |
| *Tnfa* | 5′-TGGCACCACTAGTTGGTTGTCT-3′ | 5′- AGCCTGTAGCCCACGTCGTA -3′ | S1 |
| *Il6* | 5′-ATCGTGGAAATGAGAAAAGAGTTGT-3′ | 5′- CTGCAAGTGCATCATCGTTGT -3′ | - |
| *Nphs1* | 5′- AGAAGCTCCACGGTTAGCAC -3′ | 5′- CCTGTGAAGGCTTGGCGATA -3′ | - |
| *Podxl* | 5′- TGCAACAGTCTATGGCGTCT -3′ | 5′- GGAGGTTCACAGTTTAGCTGGT -3′ | - |
| *F3* | 5′-CAATGAATTCTCGATTGATGTGG-3′ | 5′-GGAGGATGATAAAGATGGTGGC-3′ | S3 |
| *Kng2* | 5′- AGCTGTGACCTTCATCCAGGA -3′ | 5′- TGACCAAGCACCTCCTTCAG -3′ | - |
| *Eef2* | 5′- TGTCAGTCATCGCCCATGTG-3′ | 5′- CATCCTTGCGAGTGTCAGTGA -3′ | S4 |
| *S18* | 5′- AACGAACGAGACTCTGGCAT-3′ | 5′- ACGCCACTTGTCCCTCTAAG-3′ | S5 |

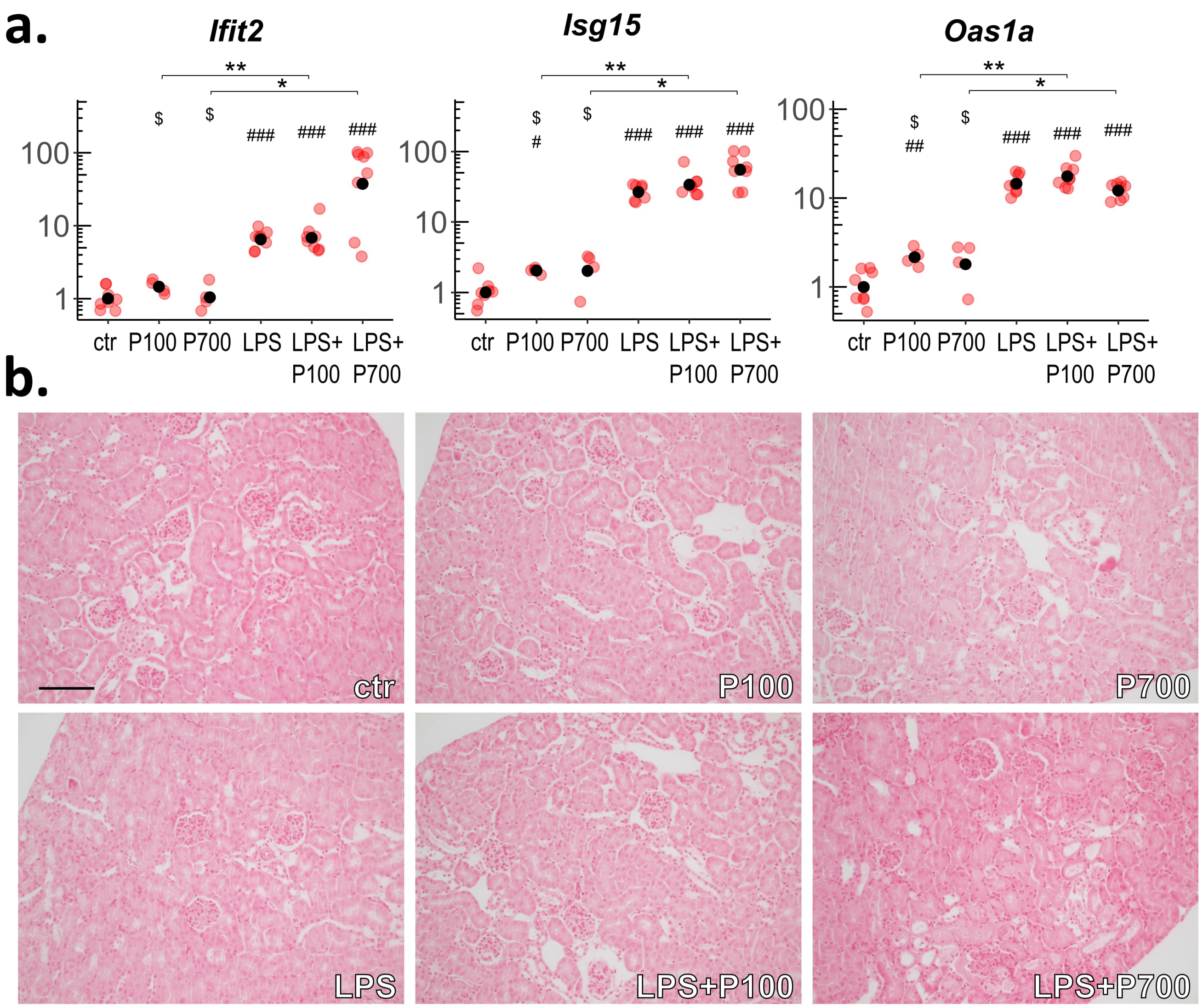


**Fig. S1. Kidney transcription of interferon stimulated genes and Von Kossa staining of kidney cortex of polyphosphate and LPS-treated mice.** (a) Transcription of interferon stimulated genes, *Ifit2*, *Isg15*, and *Oas1a*, measured by RT-qPCR from the kidney tissue of mice were treated with medium-chain polyphosphates (P100), long-chain polyphosphates (P700) alone and in combination with LPS. Red dots indicate the relative gene expression of each mouse compared to control group. Black dots indicate the geometric mean per group, scaled as one in the control group. Benjamini-Hochberg adjusted significance levels are calculated using the Mann-Whitney U test., *: $p < 0.05$, **: $p < 0.01$, ***: $p < 0.001$. Symbols used for significance: #: significance compared to the control group, $: significance compared to the LPS-treated group. (b) The Von Kossa staining showed no calcium deposits, characteristic of phosphate toxicity in the kidney tissue of mice treated with P100 or P700 alone or in combination with LPS. Scale bar 100µm.

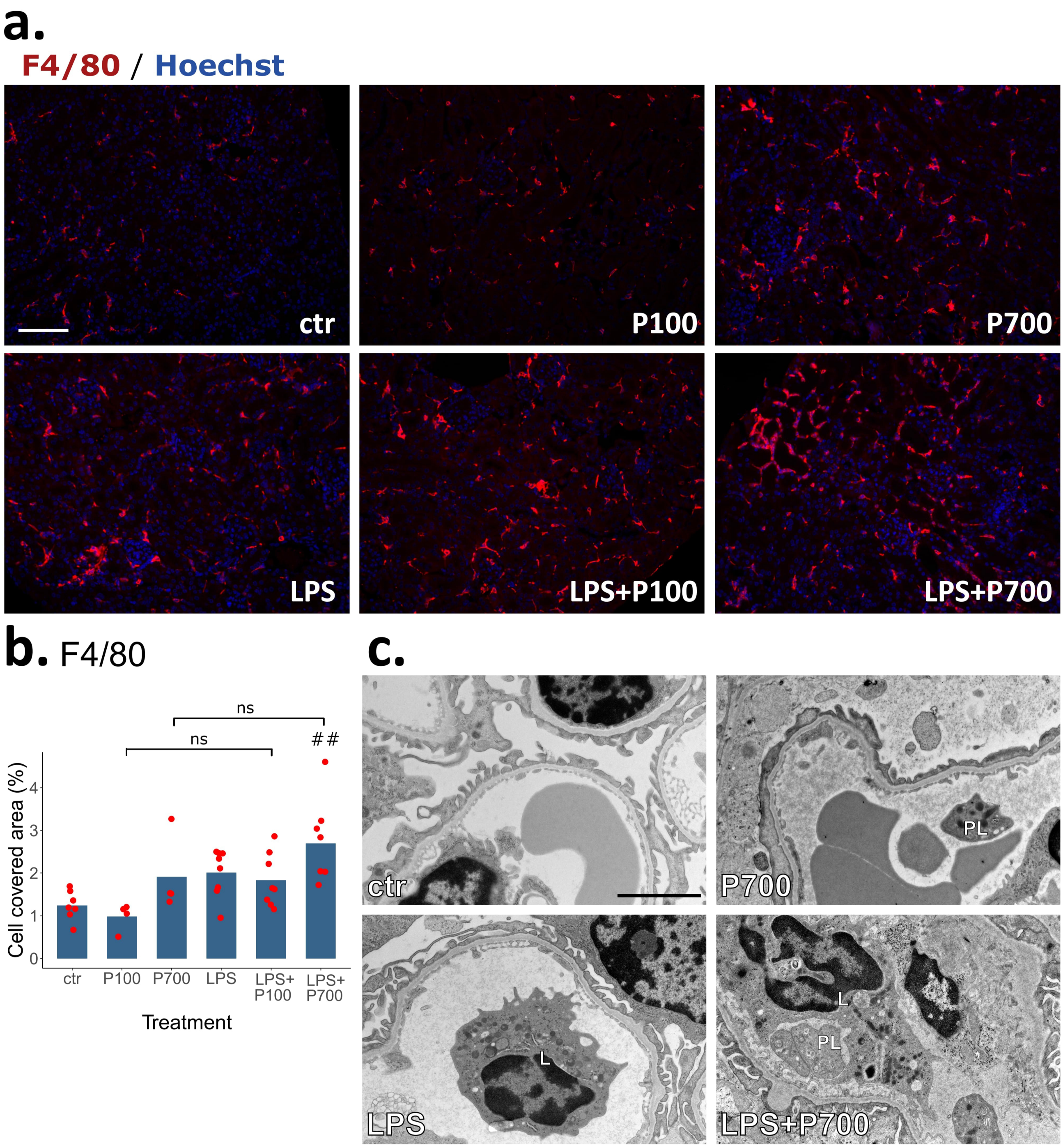


**Fig. S2. Leukocyte infiltration in the kidney cortex and glomerular capillaries of mice treated with long-chain polyphosphates and LPS.** (a) Representative images of tissue resident macrophage infiltration in the kidney cortex of mice treated with medium-chain polyphosphates (P100), long-chain polyphosphates (P700) alone and in combination with LPS, immunostained with F4/80 (red). Nuclei stained with Hoechst (blue), scale bar 100 µm. (b) Macrophage covered area in the kidney cortex. Red dots represent the mean leukocyte covered area in a frame per mouse and the bars represent the mean values per group. Benjamini-Hochberg adjusted significance levels are calculated using the pairwise Mann-Whitney U test., **: $p < 0.01$. Symbols used for significance: #: significance compared to the control group (n: 4-8 mice per group, 3-9 frames per mouse). (c) Transmission electron microscopy imaging shows intracapillary leukocytes in mice treated with LPS alone and in combination with long-chain polyphosphates (P700). 3000x magnification, scale bar 2 µm, platelets (PL), leukocytes (L).

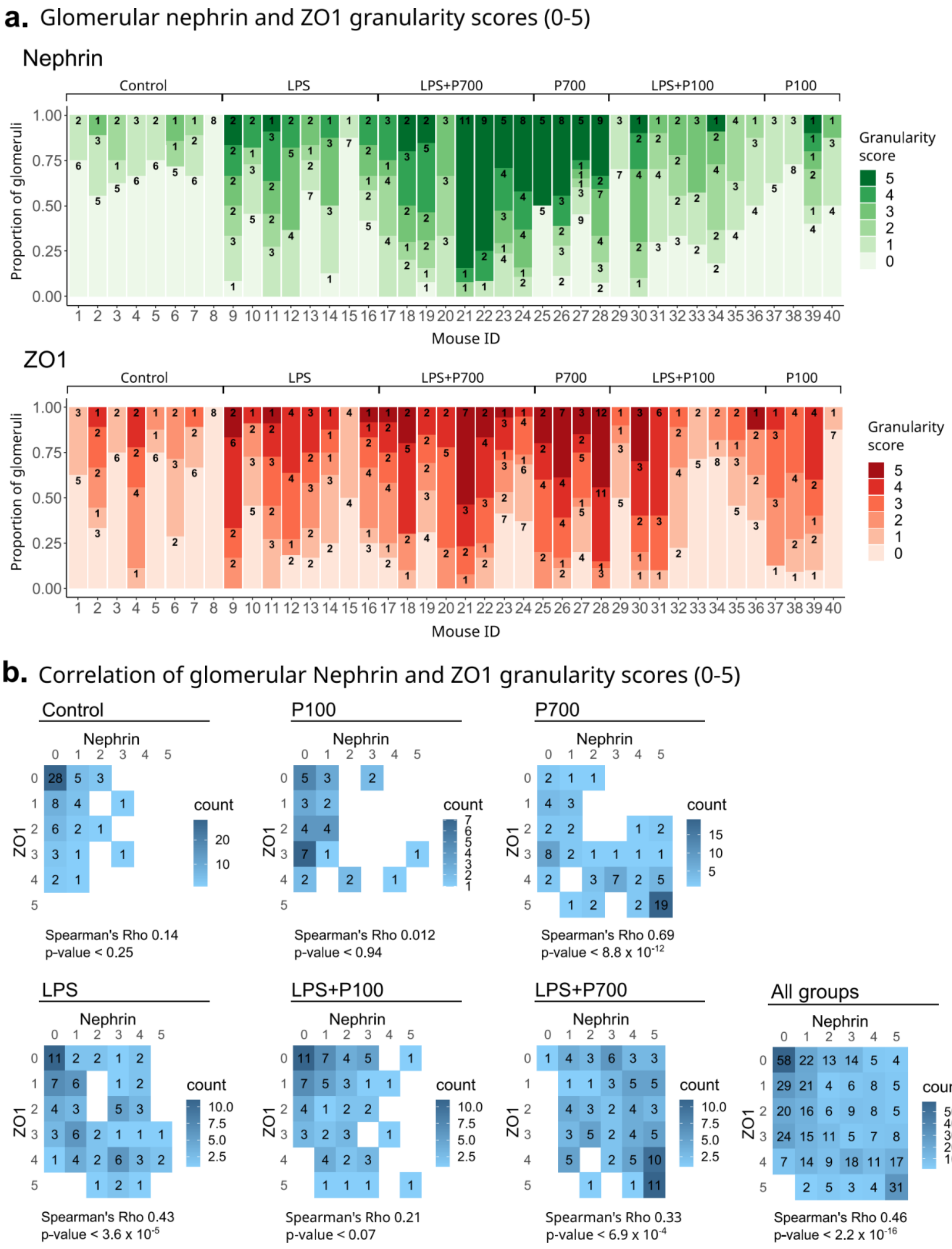

**Fig. S3. Glomerular nephrin and ZO1 granularity score proportions and correlation in polyphosphate and LPS-treated mice.** (a) Proportions of nephrin and ZO1 granularity for each mouse treated with medium-chain polyphosphates (P100), long-chain polyphosphates (P700) alone and in combination with LPS. Stacked bars represent the proportion of glomeruli affected with granularity (scored 0-5 from immunostained images) with the number of assessed glomeruli indicated inside the bars. (b) Heatmap contingency tables and spearman's correlation coefficients of glomerular nephrin and ZO1 granuarity scores (n: 4-8 mice per group, 7-27 imaged glomeruli per mouse, 444 glomeruli in total).

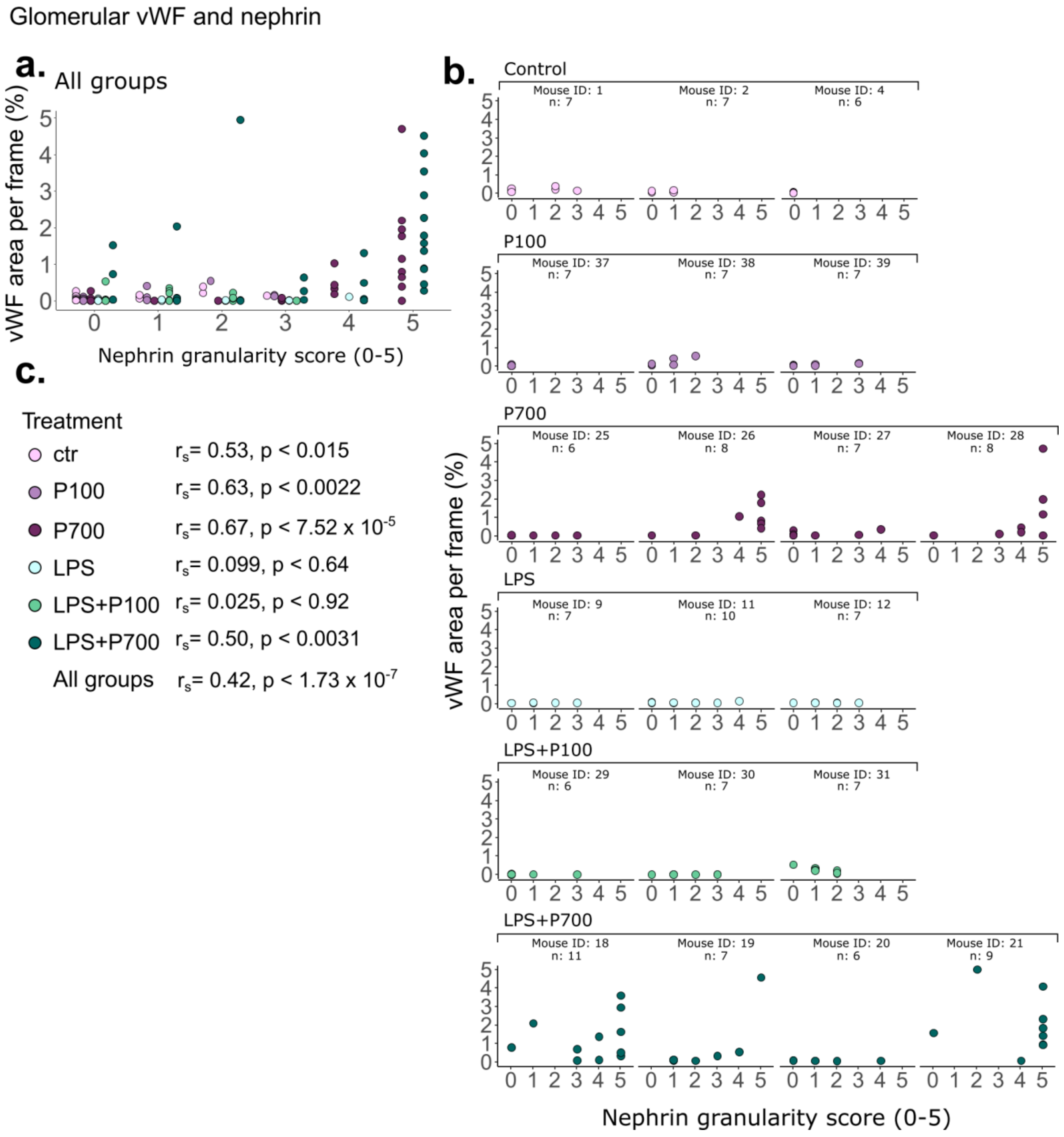


**Fig. S4. Glomerular von Willebrand factor area and nephrin granularity scores in polyphosphate and LPS-treated mice.** Quantification of von Willebrand factor-positive (vWF) area per frame and nephrin granularity score (scored 0-5 from immunostained images) for mice treated with medium-chain polyphosphates (P100), long-chain polyphosphates (P700) alone and in combination with LPS. Each dot represents one imaged glomerulus (n: 3-4 mice per group, 6-11 imaged glomeruli per mouse), (a) all groups combined, (b) number of assessed glomeruli indicated for each mouse, (c) correlation of glomerular vWF-positive area and nephrin granularity score (Spearman's correlation coefficient, $r_s$).

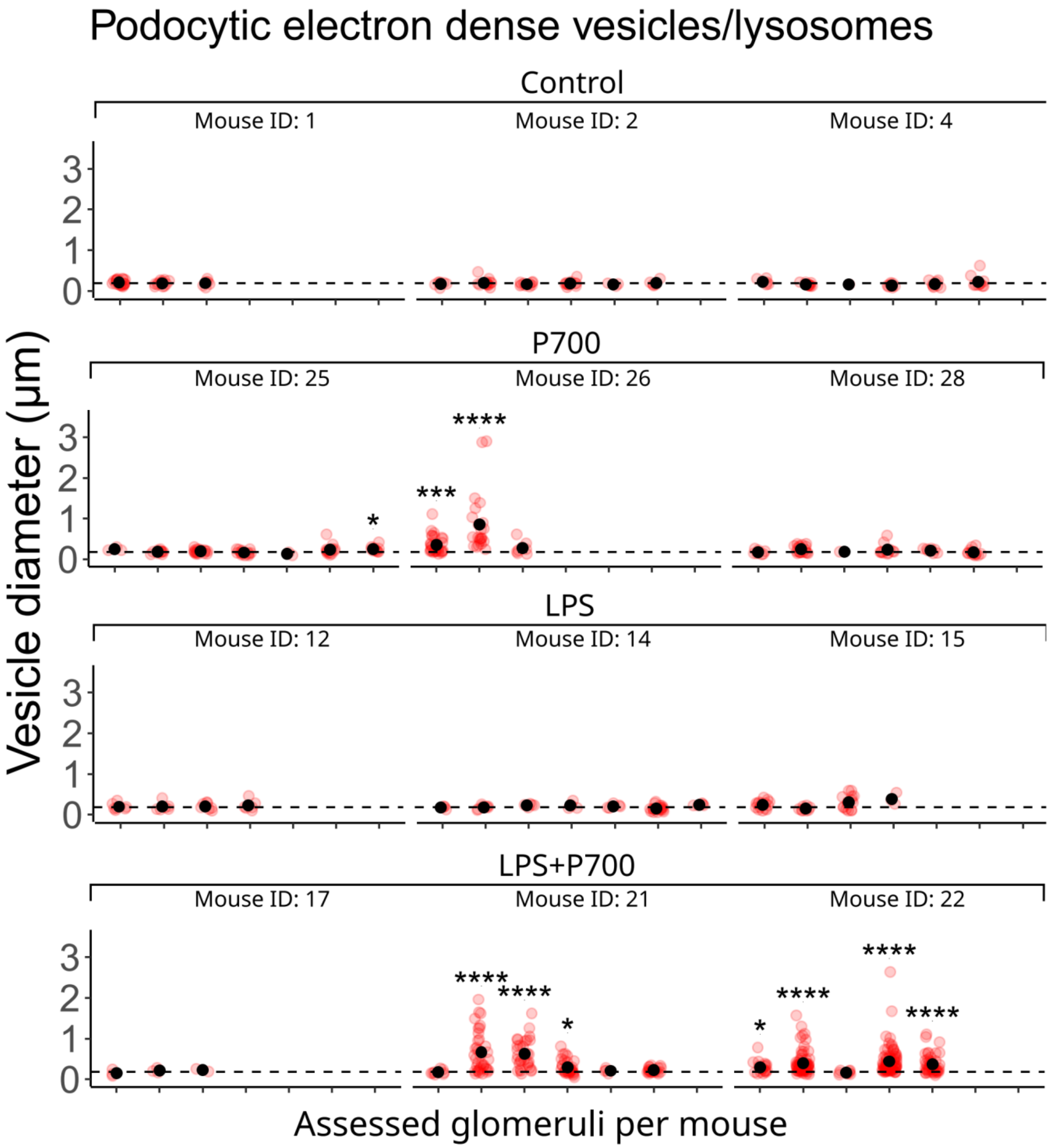


**Fig. S5. Diameter of podocytic vesicles characteristic of lysosomes, multivesicular bodies or other endosomes in transmission electron microscopy analysis**. Quantification of the diameter of podocytic vesicles characteristic of lysosomes/multivesicular bodies/endosomes in mice treated with long-chain polyphosphates (P700) and LPS. Red dots indicate individual vesicles measured in glomeruli. Black dots indicate the mean vesicle diameter per glomerulus. Benjamini-Hochberg adjusted significance levels are calculated for each glomerulus compared to the mean glomerular vesicle diameter of the healthy controls (0.18 µm, dotted line) using the Mann-Whitney U test., *: $p < 0.01$, ***: $p < 0.001$, ****: $p < 0.0001$ (n: 3 mice per group, 3-7 imaged glomeruli per mouse, 1-5 frames per glomerulus).

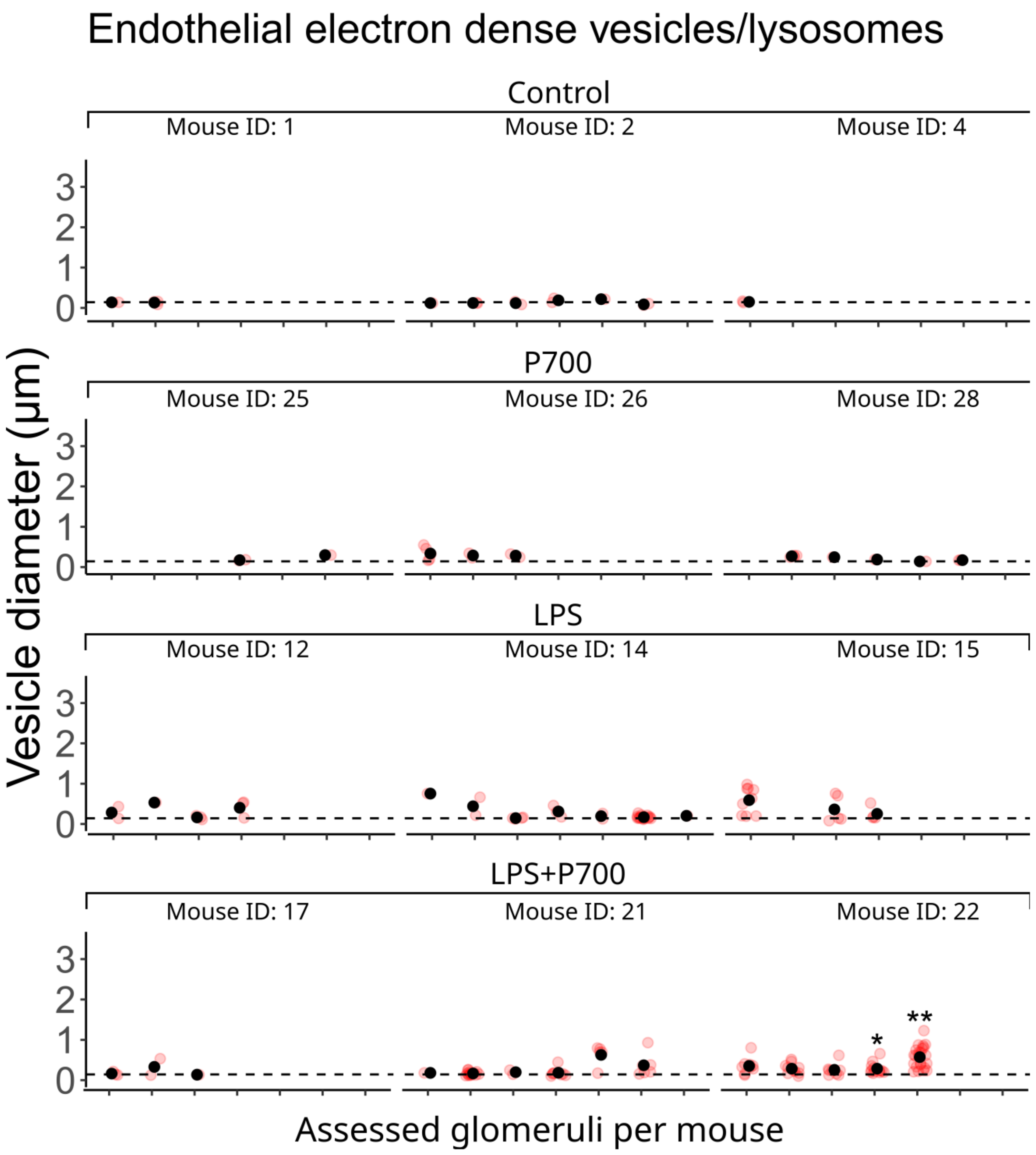


**Fig. S6. Diameter of glomerular endothelial cell vesicles characteristic of lysosomes, multivesicular bodies or other endosomes in transmission electron microscopy analysis**. Quantification of the diameter of endothelial cell vesicles characteristic of lysosomes/multivesicular bodies/endosomes in mice treated with long-chain polyphosphates (P700) and LPS. Red dots indicate individual vesicles measured in glomeruli. Black dots indicate the mean vesicle diameter per glomerulus. Benjamini-Hochberg adjusted significance levels are calculated for each glomerulus compared to the mean glomerular vesicle diameter of the healthy controls (0.14 µm, dotted line) using the Mann-Whitney U test., *: $p < 0.05$, **: $p < 0.01$ (n: 3 mice per group, 3-7 imaged glomeruli per mouse, 1-5 frames per glomerulus).
